# Structural basis of resistance to lincosamide, streptogramin A, and pleuromutilin antibiotics by ABCF ATPases in Gram-positive pathogens

**DOI:** 10.1101/2020.11.24.396648

**Authors:** Caillan Crowe-McAuliffe, Victoriia Murina, Kathryn Jane Turnbull, Marje Kasari, Merianne Mohamad, Christine Polte, Hiraku Takada, Karolis Vaitkevicius, Jörgen Johansson, Zoya Ignatova, Gemma C. Atkinson, Alex J. O’Neill, Vasili Hauryliuk, Daniel N. Wilson

## Abstract

Target protection proteins bind to antibiotic targets and confer resistance to the host organism. One class of such proteins, termed <u>a</u>ntibiotic <u>re</u>sistance (ARE) ATP binding cassette (ABC) proteins of the F-subtype (ARE ABCFs), are widely distributed throughout Gram-positive bacteria and bind the ribosome to alleviate translational inhibition by antibiotics that target the large ribosomal subunit. Using single-particle cryo-EM, we have solved the structure of ARE ABCF–ribosome complexes from three Gram-positive pathogens: *Enterococcus faecalis* LsaA, *Staphylococcus haemolyticus* VgaA_LC_ and *Listeria monocytogenes* VgaL. Supported by extensive mutagenesis analysis, these structures enable a comparative approach to understanding how these proteins mediate antibiotic resistance on the ribosome. We present evidence of mechanistically diverse allosteric relays converging on a few peptidyltransferase center (PTC) nucleotides, and propose a general model of antibiotic resistance mediated by these ARE ABCFs.

## Introduction

The bacterial ribosome is a major antibiotic target (Wilson, 2014). Despite the large size of the ribosome, and the chemical diversity of ribosome-targeting small compounds, only a few sites on the ribosome are known to be bound by clinically-used antibiotics. On the 50S large ribosomal subunit, two of the major antibiotic binding sites are the peptidyltransferase center (PTC) and the nascent peptide exit tunnel. The PTC is targeted by <u>p</u>leuromutilin, <u>s</u>treptogramin <u>A</u>, and lincosamide (PS_A_L) antibiotics, as well as phenicols and oxazolidinones (Dunkle *et al*, 2010; Matzov *et al*, 2017; Schlünzen *et al*, 2004; Tu *et al*, 2005; Wilson *et al*, 2008). Representatives of macrolide and streptogramin B classes bind at adjacent sites at the beginning of the nascent peptide exit tunnel (Dunkle *et al*., 2010; Tu *et al*., 2005).

Many mechanisms have evolved to overcome growth inhibition by such antibiotics in bacteria, among them target protection mediated by a subset of ABC family of proteins (Wilson *et al*, 2020). ATP-binding cassette (ABC) ATPases are a ubiquitous superfamily of proteins found in all domains of life, best-known as components of membrane transporters (Krishnan *et al*, 2020; Rees *et al*, 2009). A typical ABC transporter contains two nucleotide-binding domains (NBDs), each of which contribute one of two faces to an ATP-binding pocket, as well as transmembrane domains (Thomas & Tampé, 2020). Some sub-groups of ABC proteins, however, lack membrane-spanning regions and have alternative cytoplasmic functions, such as being involved in translation (Davidson *et al*, 2008; Fostier *et al*, 2020; Gerovac & Tampé, 2019). For example, in eukaryotes Rli1/ABCE1 is a ribosome splitting factor involved in recycling after translation termination, and the fungal eEF3 proteins bind the ribosome to facilitate late steps of translocation and E-site tRNA release (Andersen *et al*, 2006; Ranjan *et al*, 2020). The F-type subfamily of ABC proteins, which are present in bacteria and eukaryotes, contain at least two NBDs separated by an α-helical interdomain linker and notably lack transmembrane regions (Murina *et al*, 2019; Ousalem *et al*, 2019).

One group of bacterial ABCFs, which are termed antibiotic resistance (ARE) ABCFs (Dorrian & Kerr, 2009), confer resistance to antibiotics that bind to the 50S subunit of the bacterial ribosome (Ero *et al*, 2019; Ousalem *et al*., 2019; Sharkey & O’Neill, 2018; Wilson *et al*., 2020). Characterized ARE ABCFs are found predominantly in Gram-positive bacteria, including human and animal pathogens, typically have a restricted host specificity, and can be further divided into eight subfamilies (Allignet *et al*, 1992; Murina *et al*., 2019; Wilson *et al*., 2020). Although initially thought to act as part of efflux systems (Ross *et al*, 1990; Ross *et al*, 1989), these proteins were subsequently shown instead to bind the ribosome, oppose antibiotic binding, and to reverse antibiotic-mediated translation inhibition of translation *in vitro* (Sharkey *et al*, 2016).

Phylogenetic analyses indicate that ARE ABCFs may have arisen multiple times through convergent evolution, and that antibiotic specificity can be divergent within a related subgroup (Murina *et al*., 2019). Classified by the spectrum of conferred antibiotic resistance, ARE ABCFs can be categorized into three groups (Murina *et al*., 2019; Sharkey & O’Neill, 2018):

1. A highly polyphyletic group of ARE ABCFs that confer resistance to the PTC-binding PS_A_L antibiotics (ARE1, ARE2, ARE3, ARE5 and ARE6 subfamilies). The most well-studied representatives are VmlR, VgaA, SalA, LmrC and LsaA (Allignet *et al*., 1992; Hot *et al*, 2014; Koberska *et al*, 2020; Ohki *et al*, 2005; Singh *et al*, 2002). Additionally, a lincomycin-resistance ABCF that belongs to this group, termed Lmo0919, has been reported in *Listeria monocytogenes* (Chesneau *et al*, 2005; Dar *et al*, 2016; Duval *et al*, 2018).
2. ARE ABCFs that confer resistance to antibiotics that bind within the nascent peptide exit channel (a subset of the ARE1 subfamily, and ARE4). The most well-studied representatives are <u>M</u>acrolide and <u>s</u>treptogramin B resistance (Msr) proteins (Reynolds & Cove, 2005; Ross *et al*., 1990; Su *et al*, 2018).
3. Poorly experimentally characterized ARE ABCF belonging to subfamilies ARE7 (such as OptrA) and ARE8 (PoxtA). These resistance factors confer resistance to phenicols and oxazolidinones that bind in the PTC overlapping with the PS_A_L binding site (Antonelli *et al*, 2018; Wang *et al*, 2015; Wilson *et al*., 2020) and are spreading rapidly throughout bacteria in humans and livestock by horizontal gene transfer (Freitas *et al*, 2017; Iimura *et al*, 2020; Sadowy, 2018; Zhang *et al*, 2020).

Additionally, several largely unexplored groups of predicted novel ARE ABCFs are found in high-GC Gram-positive bacteria associated with antibiotic production (Murina *et al*., 2019).

So far, two structures of ARE ABCFs bound to the 70S ribosome have been determined (Crowe-McAuliffe *et al*, 2018; Ero *et al*., 2019; Su *et al*., 2018). In each instance, the ARE ABCF interdomain linker extended from the E-site-bound NBDs into the relevant antibiotic-binding site in the ribosome, distorting the P-site tRNA into an unusual state in the process. The tip of the interdomain linker—termed the <u>a</u>ntibiotic resistance <u>d</u>eterminant (ARD)—is not well conserved among (or sometimes even within) subfamilies, and mutations in this region can abolish activity as well as change antibiotic specificity. Mutagenesis indicates that both steric overlap between the ARD and the antibiotic, as well as allosteric reconfiguration of the rRNA and the antibiotic-binding site, may contribute to antibiotic resistance (Crowe-McAuliffe *et al*., 2018; Ero *et al*., 2019; Lenart *et al*, 2015; Su *et al*., 2018). Non-ARE ribosome-associated ABCFs that do not confer resistance to antibiotics—such as EttA—tend to have relatively short interdomain linkers that contact and stabilize the P-site tRNA (Chen *et al*, 2014). ARE ABCFs that confer resistance to PS_A_L antibiotics (such as VmlR) have extensions in the interdomain linker that allow them to reach into the antibiotic-binding site in the PTC (Chen *et al*., 2014; Crowe-McAuliffe *et al*., 2018; Lenart *et al*., 2015). The longest interdomain linkers belong to ARE ABCFs that confer resistance to macrolides and streptogramin Bs (e.g. MsrE), and such linkers can extend past the PTC into the nascent peptide exit tunnel (Su *et al*., 2018). The length of the bacterial ABCF ARD generally correlates with the spectrum of conferred antibiotic resistance. Notable exceptions to this pattern are OptrA and PoxtA ARE ABCF which have short interdomain linkers, yet still confer resistance to some PTC-binding antibiotics (Antonelli *et al*., 2018; Wang *et al*., 2015), while typically PTC-protecting ARE ABCFs such as VmlR, LsaA and VgaA, typically have comparatively long interdomain linkers (Lenart *et al*., 2015; Singh *et al*, 2001).

The available ARE ABCF-ribosome structures were generated by *in vitro* reconstitution. *Pseudomonas aeruginosa* MsrE, which confers resistance to tunnel-binding macrolides and streptogramin Bs (that inhibit translation elongation) was analyzed bound to a heterologous *Thermus thermophilus* initiation complex (Su *et al*., 2018). *Bacillus subtilis* VmlR, which confers resistance to PS_A_L antibiotics that bind in the PTC (which stall translation at initiation) was analyzed in complex with an *B. subtilis* 70S ribosome arrested during elongation by the presence of a macrolide antibiotic (Crowe-McAuliffe *et al*., 2018; Dornhelm & Högenauer, 1978; Meydan *et al*, 2019; Ohki *et al*., 2005; Orelle *et al*, 2013). Structures of native physiological complexes (such as those generated using pull-down approaches from the native host) are currently lacking.

Here we have thoroughly characterized the antibiotic resistance specificity and determined the structure of three native ARE ABCF-70S ribosome complexes using affinity chromatography and cryo-electron microscopy (cryo-EM). We selected ARE ABCFs that confer resistance to PS_A_L antibiotics in clinically-relevant Gram-positive pathogens: ARE3 representative *Enterococcus faecalis* LsaA (Singh *et al*., 2002), and ARE1 representatives *Listeria monocytogenes* Lmo0919 (Chesneau *et al*., 2005; Dar *et al*., 2016; Duval *et al*., 2018)—which we have termed VgaL—as well as the well-characterized VgaA_LC_ protein, initially isolated from *Staphylococcus haemolyticus* (Allignet *et al*., 1992; Chesneau *et al*., 2005; Jacquet *et al*, 2008; Lenart *et al*., 2015; Novotna & Janata, 2006). *Staphylococcus* and *Enterococcus* are commensal organisms that are prevalent in diverse healthcare-associated infections, and antibiotic resistance is spreading through these species (Magill *et al*, 2014; Mamtora *et al*, 2019; Mendes *et al*, 2019; Pfaller *et al*, 2019). *L. monocytogenes* is a foodborne pathogen that poses particular risk to pregnant women and immunocompromised patients (Camargo *et al*, 2016). Our structures, supported by extensive mutagenesis experiments, provide much needed insight into the mechanism by which these distinct ARE ABCFs displace antibiotics from their binding site on the ribosome to confer antibiotic resistance.

## Results

### Cryo-EM structures of native ARE ABCF-70S complexes

To obtain native ARE ABCF-70S complexes, we expressed C-terminally FLAG_3_-tagged ATPase-deficient EQ_2_ variants of *E. faecalis* LsaA, *S. aureus* VgaA_LC_, and *L. monocytogenes* VgaL in their corresponding native host bacterial species. The FLAG_3_ tag was used for affinity purification of each protein locked on the ribosomal target. The ARE ABCFs co-migrated with the 70S fraction through sucrose gradients—with the complex further stabilized in the presence of ATP in the case of LsaA and VgaA_LC_—and co-eluted with ribosomal proteins after affinity purification (Figures S1–3).

The resulting native complexes were characterized by single-particle cryo-EM (see Methods), yielding ARE–70S complexes with average resolutions of 2.9 Å for *E. faecalis* LsaA, 3.1 Å for *S. aureus* VgaA_LC_, and 2.9 Å for *L. monocytogenes* VgaL (Figure 1A-C, Table S4, Figures S4–S6). In each instance, the globular nucleotide-binding domains (NBDs) of the ARE ABCF bound in the E-site, and the α-helical interdomain linker extended towards the peptidyl-transferase center (PTC, Figure 1A–C). Additionally, a distorted tRNA occupied the P-site (Figure 1A–C), similarly to what was observed previously for *P. aeruginosa* MsrE and *B. subtilis* VmlR (Crowe-McAuliffe *et al*., 2018; Su *et al*., 2018). For the LsaA and VgaL samples, occupancy of the factor on the ribosome was high, with >95% or ∼70% of picked ribosomal particles containing LsaA or VgaL, respectively (Figures S4 and S6). By contrast, VgaA_LC_ had lower occupancy (∼60%), implying that the factor dissociated after purification and/or during grid preparation (Figure S5). *In silico* 3D classification revealed that the major class not containing VgaA_LC_ in the dataset was a 70S ribosome with P-tRNA, which could also be refined to an average resolution of 3.1 Å (Figure S5). Generally, the 50S ribosomal subunit and ARE ABCF interdomain linkers were well-resolved (Figures 1D–F and S4–S6). While ARE ABCF NBDs, occupying the E site, had a lower resolution—especially in the regions that contact the ribosomal L1 stalk and the 30S subunit—the density was nonetheless sufficient to dock and adjust homology models in each instance (Figures 1D–F and S4–S6). Densities corresponding to the 30S subunits were less clear, indicating flexibility in this region, but nonetheless sufficient to build near-complete models of each ribosome. Density corresponding to ATP and a coordinated magnesium ion was observed in both nucleotide-binding sites for each ARE ABCF (Figure 1D–F and S7). Density for the ATP bound in the peripheral nucleotide-binding site was relatively poor, with little density corresponding to the nucleobase moiety, consistent with the relaxed nucleotide specificity of these proteins (Figure S7) (Murina *et al*, 2018).

**Fig. 1.**
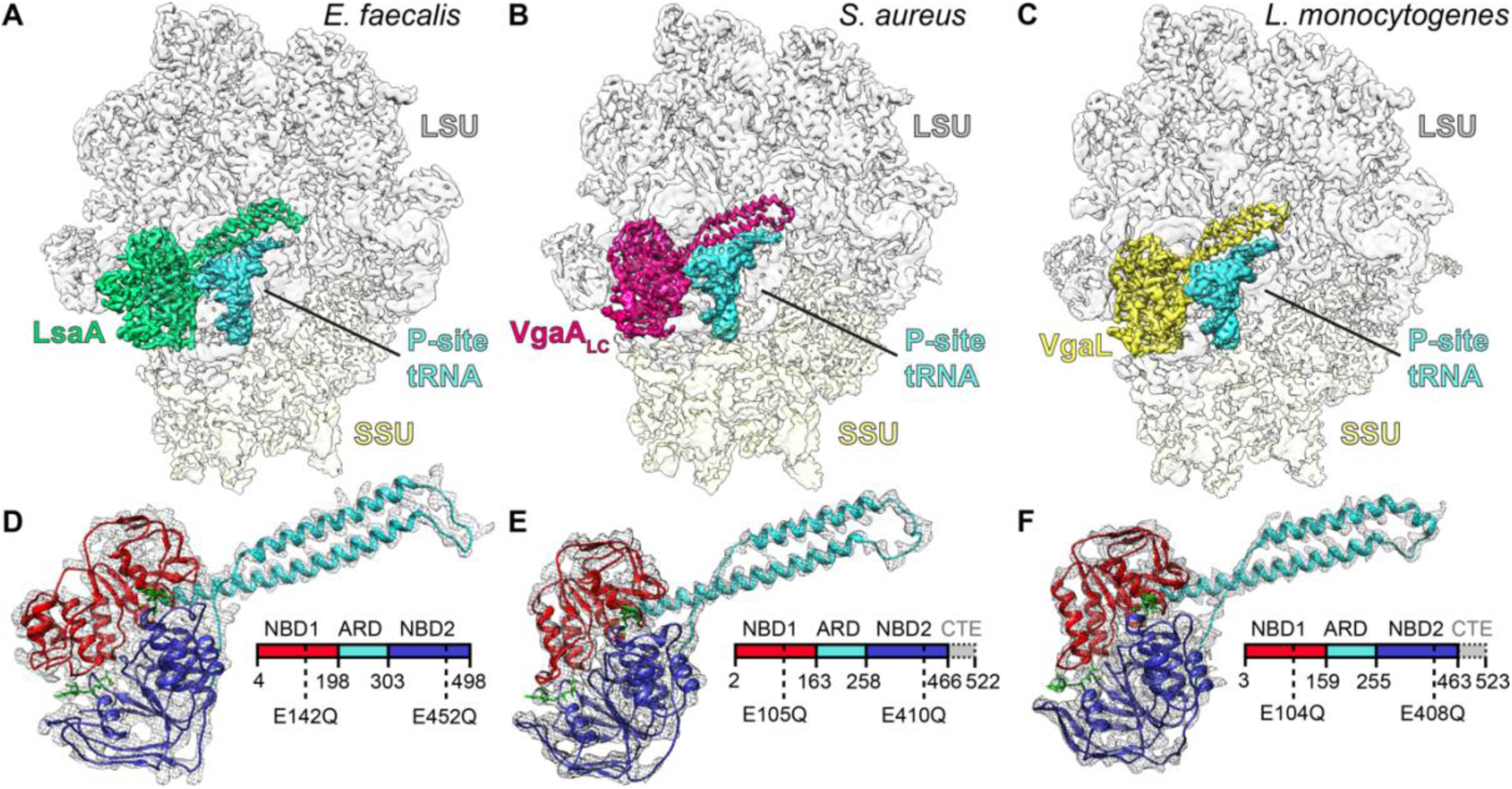
Cryo-EM structures of ARE ABCF–ribosome complexes. (**A–C**) Cryo-EM maps with isolated densities for (**A**) *E. faecalis* LsaA (green), (**B**) *S. aureus* VgaA_LC_ (magenta), (**C**) *L. monocytogenes* VgaL (yellow) as well as P-site tRNA (cyan), small subunit (SSU, yellow) and large subunit (LSU, grey). (**D–F**) Density (grey mesh) with molecular model for (**D**) LsaA, (**E**) VgaA_LC_, and (**F**) VgaL, coloured according to domain as represented in the associated schematics: nucleotide binding domain 1 (NBD1, red), antibiotic-resistance domain (ARD, cyan), nucleotide binding domain 2 (NBD2, blue) and C-terminal extension (CTE, grey, not modelled). In (**D–F**), the ATP nucleotides are coloured green.

By comparison to structures of other ABC proteins, the NBDs adopted a closed conformation bound tightly to each nucleotide (Figure S8). In each ARE ABCF–70S map, the acceptor stem of the P-site tRNA was distorted, as observed previously for MsrE and VmlR (Crowe-McAuliffe *et al*., 2018; Su *et al*., 2018). The CCA 3′ end was particularly disordered, precluding any additional density corresponding to an amino acid or nascent chain from being visualized (Figures 1A-C and S4–S6). To our knowledge, this is the first model of the ribosome from the Gram-positive pathogen *L. monocytogenes* that have been described. Additionally, we have used our high-resolution map to create an updated model of the *S. aureus* ribosome (Khusainov *et al*, 2016). Our models of the *E. faecalis* and *S. aureus* ribosomes are generally in agreement with those recently described (Golubev *et al*, 2020; Murphy *et al*, 2020).

### LsaA, Vga_LC_ and VgaL bind to translation initiation states

In each cryo-EM map, the P-site tRNA body was sufficiently well-resolved so as to unambiguously assign the density to initiator tRNA^fMet^, on the basis of (i) general fit between sequence and density, (ii) the well-resolved codon-anticodon interaction, and (iii) a characteristic stretch of G:C base pairs found in the anticodon stem loop of tRNA^fMet^ (Figure 2A–C). Additionally, in the small subunit mRNA exit tunnel, density corresponding to a putative Shine-Dalgarno–anti-Shine-Dalgarno helix was observed, consistent with the ARE ABCF binding to an initiation complex containing tRNA^fMet^ (Figure 2D). LsaA–*E. faecalis* 70S samples were further analyzed with a custom tRNA microarray, which confirmed tRNA^fMet^ was the dominant species found in the sample (Figure 2E). Collectively, these observations indicate that in our structures the majority of the ARE ABCFs are bound to 70S translation initiation complexes.

**Figure 2.**
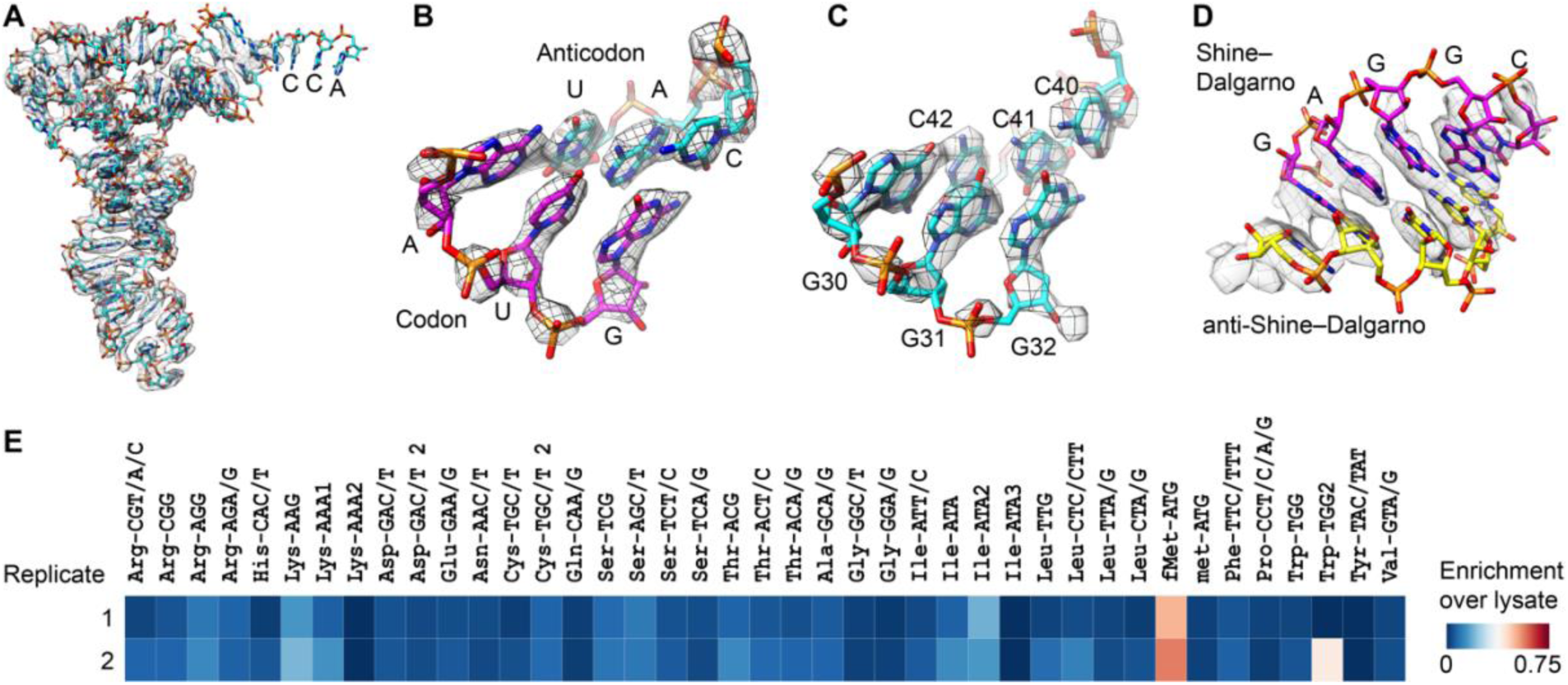
The LsaA–70S complex contains an initiator tRNA and SD-helix. (**A–D**) Isolated density (grey mesh) with molecular models (sticks) for (**A**) initiator tRNA^fMet^ (cyan), interaction between AUG start codon of the mRNA (magenta) and anticodon of initiator tRNA^fMet^ (cyan) in the P-site, (**C**) three G-C base pairs specific to the initiator tRNA^fMet^ (cyan), and (**D**) helix formed between Shine-Dalgarno (SD) sequence of the mRNA (magenta) and anti-SD of the 16S rRNA (yellow). (**E**) Replicate tRNA microarray analysis of the LsaA–70S complex, illustrating the enrichment of initiator tRNA^fMet^ in the LsaA–70S complex over the lysate. Confidence intervals between replicates were 92%.

Further examination of the LsaA–70S volume revealed weak density in the ribosomal A site (Figure S4F), suggesting that some complexes had entered into the first elongation cycle. This was unexpected, as the distorted P-site tRNA is predicted to overlap with an accommodated A-site tRNA, although as noted would be compatible with a pre-accommodated A/T-tRNA (Crowe-McAuliffe *et al*., 2018). A mask around the A site was used for partial signal subtraction, and focused 3D classification was used to further sub-sort the LsaA–70S volume. One class, containing approximately one third of the particles, was shown to indeed contain a tRNA in the A site (Figures S4, S9A). This tRNA was poorly resolved, suggesting flexibility, and was slightly rotated compared to a canonical, fully accommodated A-site tRNA, and, as for the P-site tRNA, the acceptor stem was significantly disordered and displaced (Figure S9B,C). This state likely reflects an incomplete or late-intermediate accommodation event, as observed previously when translation is inhibited by PTC binding antibiotics hygromycin A or A201A, both of which were shown to sterically exclude the acceptor stem of a canonical A-site tRNA (Polikanov *et al*, 2015). A very weak density corresponding to an A-site tRNA was also observed in VgaA_LC_ and VgaL volumes, but sub-classification was unsuccessful for these datasets.

VgaA_LC_ and VgaL, both of which belong to the ARE1 subfamily—although not LsaA, which belongs to the ARE3 subfamily—contain a short C-terminal extension predicted to form two α-helices (Crowe-McAuliffe *et al*., 2018; Murina *et al*., 2019). Although not conserved among all AREs, deletion of the CTE abolished antibiotic resistance in VmlR and reduces antibiotic resistance in VgaA, implying that this extension is necessary for function in some ARE ABCFs (Crowe-McAuliffe *et al*., 2018; Jacquet *et al*., 2008). Density for this region, which emanates from NBD2 and was located between ribosomal proteins uS7 and uS11, was present in the VgaA_LC_–70S and VgaL–70S maps and was essentially consistent with the position of the VmlR C-terminal extension, although was not sufficiently resolved to create a model for this region. Although bound close to the mRNA exit channel, the CTEs of VgaA_LC_ and VgaL did not contact the Shine-Dalgarno–anti-Sine-Dalgarno helix of the initiation complexes, indicating they are not critical for substrate recognition in these ARE ABCFs (Figure S10).

### The location and conformation of short and long ARDs on the ribosome

The ARD loop, positioned between the two long α-helices that link the NBDs, is a critical determinant of antibiotic resistance (Crowe-McAuliffe *et al*., 2018; Lenart *et al*., 2015; Murina *et al*., 2018; Sharkey *et al*., 2016; Su *et al*., 2018). Despite sharing a similar antibiotic specificity profile, the ARDs of LsaA, VgaA_LC_, VgaL, and VmlR are divergent in both amino acid composition and length, which is consistent with the polyphyletic nature of this group but precludes confident sequence alignment of this region (Figure 3A). Despite such sequence divergence, the position of the ARDs on the ribosome is broadly similar in each instance (Figure 3B–G). By comparison to tiamulin, which overlaps with the aminoacyl moieties of A- and P-tRNAs in the PTC, VmlR, LsaA, VgaA_LC_, and VgaL are all positioned similarly on the ribosome, with the ARD backbone adjacent to the antibiotic binding site (Figure 3B–F) (Polikanov *et al*., 2015; Schlünzen *et al*., 2004). Compared to VmlR, the additional residues in the ARDs of LsaA, VgaA_LC_, and VgaL extend away from the antibiotic binding site, towards the CCA 3′ end of the distorted P-tRNA (Figure 3C–F). By contrast, MsrE, which confers resistance to tunnel-binding antibiotics deeper in the ribosome, has a longer ARD that extends both past the PTC to approach the macrolide/streptogramin A binding site, as well as towards the distorted P-tRNA (Figure 3A, G) (Dunkle *et al*., 2010; Su *et al*., 2018). Thus, the length of the ARD does not necessarily provide insights into the extent to which the ARD will extend into the ribosomal tunnel and thus one cannot easily predict whether long ARDs will confer resistance to macrolide antibiotics.

**Fig. 3.**
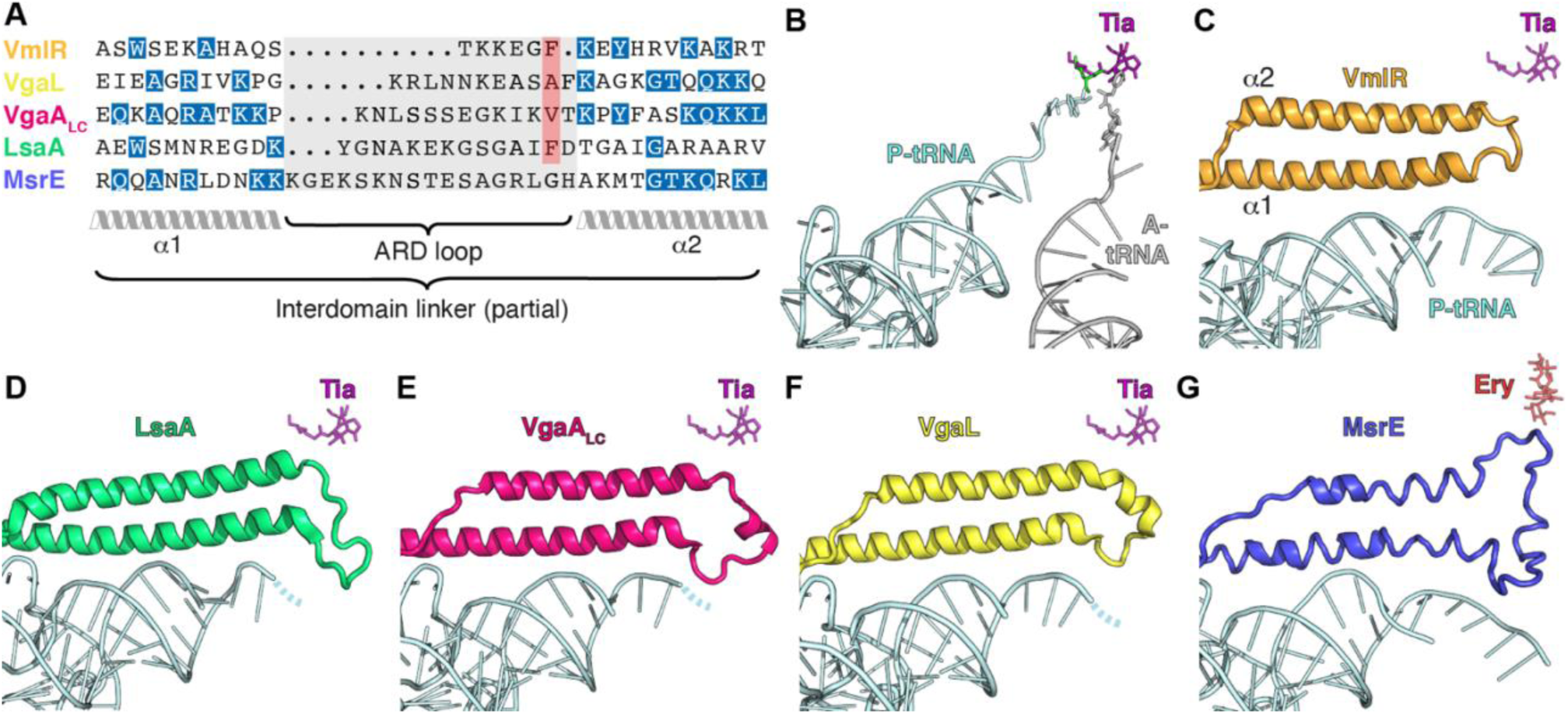
Comparison of the ARD loops of different ARE ABCFs. (**A**) The sequence length of the ARD loops differs significantly for VmlR, VgaL, VgaA_LC_, LsaA and MsrE. Although the lack of sequence homology precludes accurate sequence alignment of the ARD loops, the red highlighted residues can be aligned structurally. (**B–G**) Comparison of the positions of (**B**) A-site tRNA (grey) and P-site tRNA (cyan) from pre-attack state (PDB 1VY4) (Polikanov *et al*, 2014), with shifted P-site tRNA (cyan) and ABCF ARD from ribosome complexes containing (B) VmlR (orange, PDB 6HA8) (Crowe-McAuliffe *et al*., 2018), (**D**) LsaA (green), (**E**) VgaA_LC_ (magenta), (**F**) VgaL (yellow), and (**G**) MsrE (blue, PDB 5ZLU) (Su *et al*., 2018). In (**B–G**), the relative position of either tiamulin (Tia, magenta, PDB 1XBP) (Schlünzen *et al*., 2004) or erythromycin (Ery, red, PDB 4V7U) (Dunkle *et al*., 2010) has been superimposed.

### Position of the ARDs with respect to PS_A_L antibiotic binding site

We next made a careful comparison of the LsaA, VgaA_LC_, and VgaL ARDs with the binding sites of relevant antibiotics within the PTC (Figure 4A, B) (Dunkle *et al*., 2010; Matzov *et al*., 2017; Schlünzen *et al*., 2004; Tu *et al*., 2005). For LsaA, the side chain of Phe257 overlapped with the binding sites of tiamulin, virginiamycin M, and lincomycin, but was not close to erythromycin (Figure 4A–C), consistent with the spectrum of antibiotic resistance conferred by this protein (Table S1). In the VgaA_LC_ ARD, Val219 was situated close to tiamulin and virginiamycin M, and had a modest predicted overlap with lincomycin (Figure 4D). Notably, in the closely related variant VgaA, which has a similar specificity with modestly higher resistance to tiamulin and virginiamycin M, residue 219 is a glycine, which we predict would not overlap with the PS_A_L binding site (Lenart *et al*., 2015). Thus, VgaA_LC_ confers resistance to virginiamycin M and tiamulin despite the lack of overlap between the ARE ABCF and the antibiotic binding site (Table S2). For VgaL, the closest residue to the PS_A_L binding site was Ala216, which had no predicted overlap with tiamulin, virginiamycin M, or lincomycin (Figure 4E). Strikingly, VgaL therefore confers resistance to lincomycin, virginiamycin M, and tiamulin without directly overlapping the binding sites of these antibiotics. In summary, there was no general pattern of overlap or non-overlap with the PS_A_L binding sites among LsaA, VgaA_LC_, and VgaL.

**Fig. 4.**
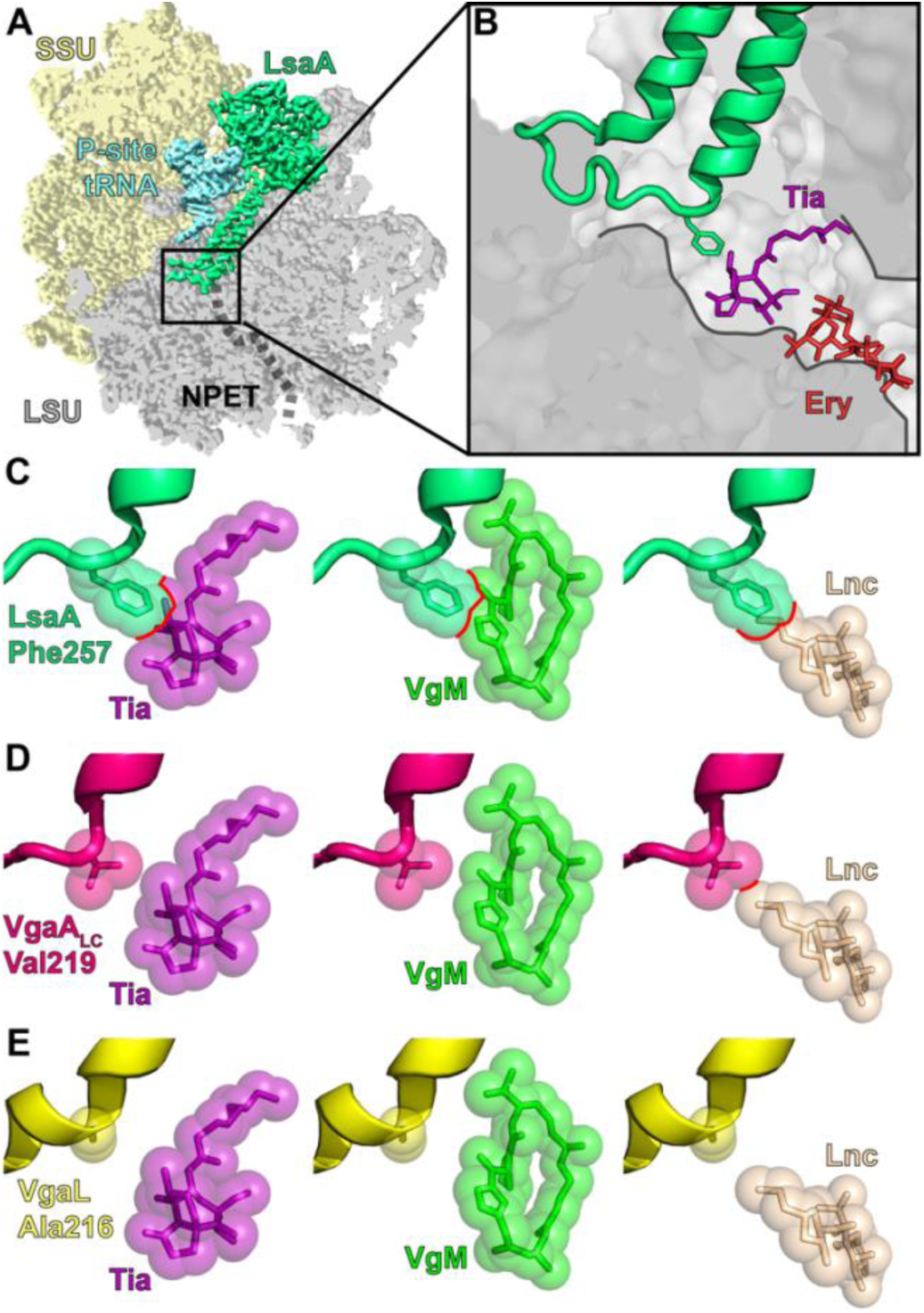
Interaction of LsaA, VgaA_LC_ and VgaL at the peptidyltransferase centre. (**A–B**) LsaA and distorted P-site tRNA superimposed on a transverse section of the large subunit (LSU, grey) to reveal (**A**) the ARD of LsaA extending into the nascent polypeptide exit tunnel (NPET) and (**B**) the relative position of Phe257 of LsaA to tiamulin (Tia, purple, PDB 1XBP) (Schlünzen *et al*., 2004) and erythromycin (Ery, red, PDB 4V7U) (Dunkle *et al*., 2010). (**C–E**) Relative position of LsaA (green, row **C**), VgaA_LC_ (pink, row **D**) and VgaL (yellow, row **E**) to tiamulin (Tia, purple, PDB 1XBP), virginiamycin M (VgM, lime, PDB 1YIT) (Tu *et al*., 2005), lincomycin (Lnc, tan, PDB 5HKV) (Matzov *et al*., 2017). Clashes in **C–E** are shown with red outlines.

### Mutational analysis of LsaA and VgaA_LC_ ARDs

Our models of the ARD loops allowed us to design and test mutants for capacity to confer antibiotic resistance. When LsaA Phe257, which directly overlaps the PS_A_L binding site (Figure 4C), was mutated to alanine, no change in resistance was observed (Figure S11). By contrast, mutation of Lys244, which is not situated close to the PS_A_L binding sites but forms a hydrogen bond with 23S rRNA G2251 and G2252 of the P-loop (*Escherichia coli* numbering is used for 23S rRNA nucleotides), nearly abolished antibiotic resistance activity (Figure S11 and S12A–C). Combined, these observations indicate that LsaA does not confer resistance *via* simple steric occlusion, and that interactions with the P-loop may be required for positioning the LsaA ARD. For VgaA_LC_, extensive alanine mutations within the ARD were explored (Table S2). As expected from the above analyses and natural variants, mutating Val219—the only residue in VgaA_LC_ that sterically overlaps the LS_A_P binding site—did not affect the antibiotic resistance profile. Three residues at the beginning of α2, directly after the ARD loop, were required for resistance: Tyr223, which stacks with U2585 (part of the pleuromutilin and lincomycin binding sites); Phe224, which stacks with A2602 held in the center of the ARD; and Lys227, which forms a hydrogen bond with the 5′ phosphate of C2601 (Table S2). These residues do not overlap with the PS_A_L binding site, but may be required to position the ARD in the PTC to impede antibiotic binding, or for the folding of the ARD itself (Figure S12D–F). In the naturally variable VgaA_LC_ ARD, mutation of Ser213, which sits adjacent to U2506 and C2507 (Figure S12E), to alanine similarly reduced antibiotic resistance (Table S2). Of note, mutating the most conserved residue among VgaA variants in this region, Lys218, did not substantially affect resistance (Table S2) (Vimberg *et al*, 2020). Extensive alanine substitutions in the surrounding residues that contact the 23S rRNA (Figure S12D–F) either did not affect, or had only a mild influence on, the antibiotic resistance conferred by this protein (Table S2). In summary, mutation of VgaA_LC_ residues that interact with 23S rRNA nucleotides that form part of the LS_A_P binding pocket affected antibiotic-resistance activity.

### Modulation of the ribosomal antibiotic binding site by ARE ABCFs

We next sought to explore how the ARDs of LsaA, VgaA_LC_, and VgaL affect the conformation of the ribosomal PTC. The 23S rRNA A2602, which is flexible in the absence of tRNAs and positioned between the P- and A-tRNAs during peptidyl transfer, is bound and stabilized by all structurally characterized ARE ABCFs. In LsaA and VmlR, a tryptophan stacks and stabilizes A2602 in a flipped position (Figure S13) (Crowe-McAuliffe *et al*., 2018). In VgaA_LC_, VgaL, and MsrE, A2602 is instead positioned within the ARD loop, interacting with multiple residues from the ARE (Figure S13) (Su *et al*., 2018). We have labelled five regions of domain V of the 23S rRNA, which form the PTC, PTC loops (PLs) 1–5 (Figure 5A) (Polacek & Mankin, 2005). There was a significant overlap between nucleotides that form the PS_A_L binding pockets, nucleotides that were shifted when LsaA, VgaA_LC_, or VgaL bound the 70S, and nucleotides known to be mutated or modified in antibiotic-resistant strains of bacteria (summarized in Figure 5A). Broadly, changes to the PTC were similar between the VgaA_LC_- and VgaL-bound 70S structures, consistent with the grouping of these proteins together in the ARE1 subfamily (Figures 5E–G, S14, S15) (Murina *et al*., 2019). Loop PL3, which contains nucleotides A2503 to U2506, was shifted upon binding of each ARE ABCF (Figure 5B–G). However, no residues from VgaA_LC_ or VgaL directly contact PL3 (Figures 5E–G, S14 and S15). Rather, these ARE ABCFs directly displace PL2, which ordinarily positions PL3, perhaps thereby facilitating the distorted conformation of PL3 that is incompatible with antibiotic binding. In the VgaA_LC_-bound state, U2585, which was poorly ordered in the LsaA- and VgaL-bound 70S, stacks with Tyr223 and would not be available to interact with tiamulin or virginiamycin M (Figure S14D–F). Substituting VgaA_LC_ Tyr223 to alanine diminished antibiotic resistance, indicating that the reposition of U2585 contributes to antibiotic resistance conferred by this ARE ABCF (Table S2 and Figure S12F). 23S rRNA U2506 is additionally displaced in the VgaA_LC_- and VgaL-bound 70S compared to the tiamulin- or lincomycin-bound 70S, potentially disrupting the binding site of these antibiotics (Figures S14A–C, S15A–C). By contrast, LsaA induced the most dramatic rearrangements in the PTC, with U2504 and G2505 in the LsaA-bound state predicted to strongly clash with each relevant antibiotic bound to the ribosome (Figure S5E–G and S16A–C). In the LsaA-bound ribosome, A2453 is shifted slightly away from the PTC and pairs with G2499 instead of U2500. This allows C2452, which normally pairs with U2504 to form part of the PS_A_L binding pocket, to instead hydrogen-bond with U2500, thereby freeing U2504, and PL3 more generally, to reposition when LsaA is bound (Figure S16D, E).

**Fig. 5.**
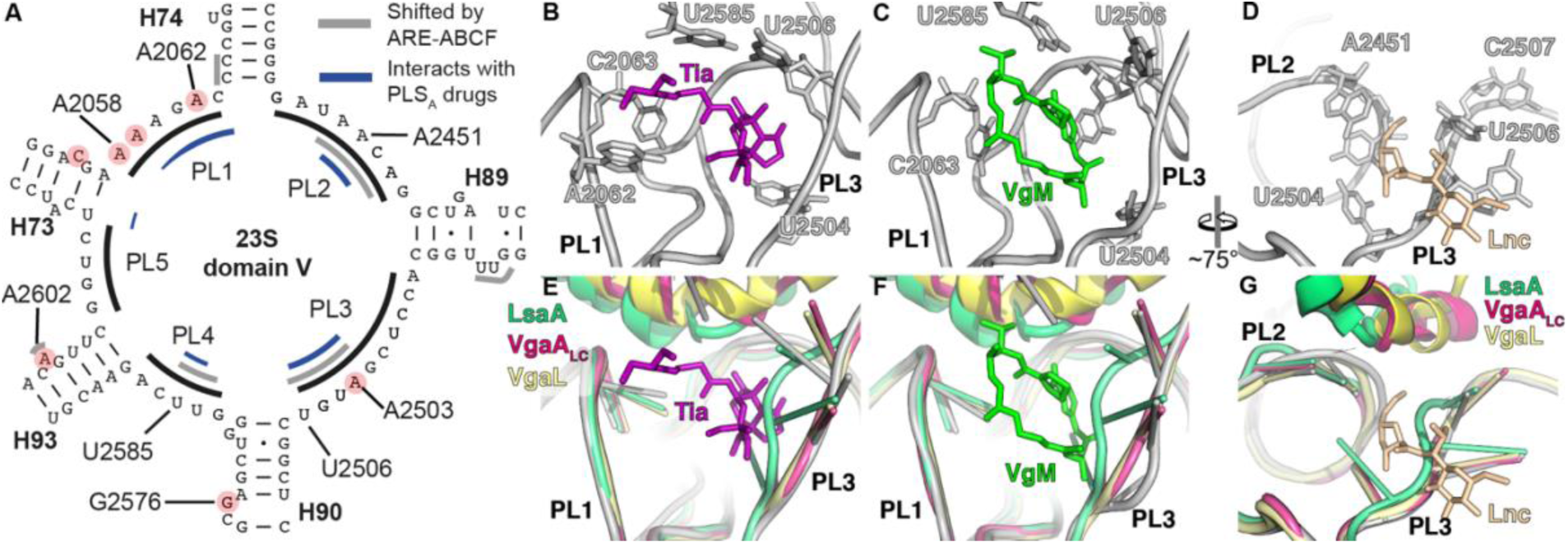
ARE ABCF binding induces allosteric conformational changes at the PTC. (**A**) Secondary structure of peptidyltransferase ring within domain V of the 23S rRNA, highlighting residues within PTC loops 1**–**4 (PL1**–**4) that (i) comprise the binding site of PS_A_L antibiotics (blue), (ii) undergo conformational changes upon ARE ABCF binding (grey) and (iii) confer resistance to PS_A_L antibiotics (red circles). (**B–D**) Binding site of (**B**) tiamulin (Tia, magenta, PDB 1XBP; (Schlünzen *et al*., 2004), (**C**) virginiamycin M (VgM, lime, PDB YIT; (Tu *et al*., 2005) and (**D**) lincomycin (Lnc, tan, PDB 5HKV) (Matzov *et al*., 2017) on the ribosome. (**E–G**) Comparison of conformations of rRNA nucleotides comprising the (**E**) Tia, (**F**) VgM and (**G**) Lnc binding site (shown as grey cartoon ladder representation), with rRNA conformations when LsaA (green), VgaA_LC_ (magenta) or VgaL (yellow) are bound.

## Discussion

### Model of antibiotic resistance mediated by LsaA, VgaA_LC_, and VgaL

These observations allow us to propose a model for how ARE ABCFs confer antibiotic resistance to the host organism (Figure 6). PS_A_L antibiotics have binding sites overlapping with the nascent polypeptide chain, and inhibit translation at, or soon after, initiation (Figure 6A) (Dornhelm & Högenauer, 1978; Meydan *et al*., 2019; Orelle *et al*., 2013). The incoming ARE ABCF binds in the E-site, triggering closure of the L1 stalk and inducing a distorted conformation of the P-tRNA. The ARD disrupts the antibiotic binding pocket in the PTC, causing drug release (Figure 6B). An incoming ternary complex delivers a tRNA to the A-site, which upon ARE ABCF egress and successful accommodation ‘sweeps’ the 3′ end of the P-tRNA into the PTC (Figure 6C, D). The trigger for nucleotide hydrolysis and exit of the ARE ABCF from the E site is unknown. We propose that rapid peptidyl transfer then creates a short nascent chain that overlaps with the antibiotic binding site, preventing re-binding of the PS_A_L drug until the next round of translation (Figure 6D). Alternatively, an A-tRNA may partially accommodate on the stalled initiation complex prior to ARE ABCF binding, and become distorted as part of a ‘knock-on’ effect of P-tRNA disruption, consistent with the ability of ARE ABCFs to ‘reset’ the P-tRNA independently of additional accommodation events (Murina *et al*., 2018). In this model, potentially only one round of ATP hydrolysis per translation cycle is necessary to confer resistance.

**Fig. 6.**
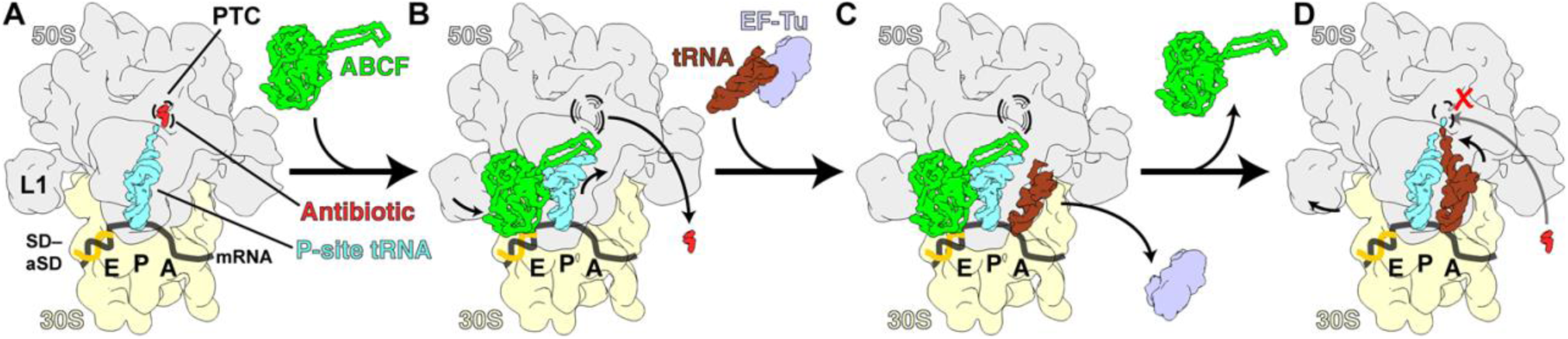
Model for ribosome protection by ARE ABCFs VmlR, LsaA, VgaA_LC_ and VgaL. **(A)** PS_A_L-stalled ribosomes containing an initiator-tRNA in the P-site are recognized by the ARE ABCFs such as VmlR, LsaA, VgaA_LC_ and VgaL, which bind to the E-site of the ribosome with a closed ATP-bound conformation. (**B**) Binding of the ARE ABCF induces a shifted P-site tRNA conformation in the ribosome allowing the ARD of the ARE ABCF to access the peptidyl-transferase center (PTC). The ARD induces conformational changes within the 23S rRNA at the PTC that promotes dissociation of the drug from its binding site (shown as dashed lines). (**C**) aminoacyl-tRNAs can still bind to the ARE ABCF-bound ribosomal complex, but cannot accommodate at the PTC due to the presence of the ABCF and shifted P-site tRNA conformation. (**D**) Hydrolysis of ATP to ADP leads to dissociation of ARE ABCF from the ribosome, which may allow the peptidyl-tRNA as well as the incoming aminoacyl-tRNA to simultaneously accommodate at the PTC. Peptide bond formation can then ensue, converting the ribosome from an initiation to an elongation (pre-translocation) state, which is resistant to the action of initiation inhibitors, such as PS_A_L antibiotics.

ARE ABCFs such as LsaA, VgaA_LC_, VgaL, and VmlR confer resistance to PS_A_L antibiotics but not phenicols or oxazolidinones (Sharkey & O’Neill, 2018). This observation has been puzzling, as both groups of antibiotics have overlapping binding sites (Dunkle *et al*., 2010; Matzov *et al*., 2017; Schlünzen *et al*., 2004; Tu *et al*., 2005; Wilson *et al*., 2008). However, phenicols and oxazolidinones inhibit translation during elongation at specific motifs (Marks *et al*, 2016; Orelle *et al*., 2013), while PS_A_L antibiotics instead inhibit translation at the initiation stage (Dornhelm & Högenauer, 1978; Meydan *et al*., 2019; Orelle *et al*., 2013). The apparent specificity of LsaA, VgaA_LC_, and VgaL for initiation complexes in our immunoprecipitations (Figure 2) matches the specificity of the antibiotics to which they confer resistance.

Do EQ_2_-substituted ATPase-deficient variants of ARE ABCF, like the ones used in this study, bind the ribosome in the pre- or post-antibiotic-dissociation state (Figure 6B)? Although direct evidence is lacking, three reasons lead us to propose that these proteins are bound in the post-antibiotic-release state:

1. In the case of VgaA_LC_ and VmlR the position of the ARD directly overlaps with the antibiotic binding site. Although the side chain of the overlapping amino acid is not critical for antibiotic resistance in most instances, the overlap nonetheless implies mutually exclusive binding.
2. MsrE-EQ_2_ stimulates dissociation of azithromycin from the ribosome (Su *et al*., 2018).
3. Our attempts to form complexes containing both antibiotic and ARE ABCF have been unsuccessful, resulting in exclusive binding of either the ARE ABCF or the antibiotic, similarly to what we observed for TetM, a tetracycline-resistance ribosome protection protein (Arenz *et al*, 2015).

How does the ARE ABCF ARD mediate antibiotic resistance (Figure 6B, C)? In one model, by analogy to the TetM tetracycline resistance protein (Arenz *et al*., 2015; Wilson *et al*., 2020), the ARD may induce antibiotic dissociation by a direct steric overlap with the antibiotic. In the case of VmlR, substitutions of the Phe237 residue that overlaps the binding site of PS_A_L antibiotics affect resistance to one of three relevant antibiotics, indicating that both direct steric overlap and an indirect, allosteric mechanism can contribute to resistance (Crowe-McAuliffe *et al*., 2018). In the case of MsrE substitution of Leu242, which overlaps with the erythromycin binding site, as well as adjacent residues abolished or severely reduced the antibiotic resistance activity of this protein (Su *et al*., 2018). In both cases, a mixture of direct steric overlap and allostery is consistent with the available data (Ero *et al*., 2019). The ARDs of LsaA, VgaA_LC_, and VgaL either do not directly overlap with the PS_A_L binding site, or where there is an overlap, as with LsaA Phe257 and VgaA_LC_ Val219, the side chains are not essential for resistance, implicating an allosteric mechanism for these proteins (Figures 4–5, S11, S12, Table S2). Alanine mutagenesis instead indicates that the side chains of residues surrounding the amino acid closest to the antibiotic-binding pocket, as well as those that contact the 23S rRNA, are necessary for resistance (Figures S11, S12 and Table S2). These residues may position the ARD in the PTC. No single set of 23S rRNA rearrangements was identical among LsaA, VgaA_LC_, and VgaL, although displacement of PTC loop PL3, especially residue U2504, was ultimately observed in each ARE ABCF–70S structure (Figure 5).

In summary, we present three new structures of ARE ABCFs bound to 70S ribosomes from relevant Gram-positive pathogenic bacteria and present the first model of the ribosome from *Listeria monocytogenes*. Our structures and mutagenesis experiments support an allosteric mechanism of ARE ABCF action, and hint at a rationalization for the specificity of LsaA, VgaA_LC_, and VgaL for PS_A_L antibiotics. Each ARE ABCF binds the 70S similarly as observed for other bacterial ABCF proteins, but alters the geometry of the PTC distinctively, consistent with the convergent evolution—and divergent sequences—of this class of ABCF proteins.

## Acknowledgments

We are grateful to Barbara E. Murray for sharing *E. faecalis* Δ*lsaA* (*lsa*::Kan) strain TX5332 (Singh *et al*., 2002), Jose A. Lemos for sharing pCIE_cam_ plasmid vector (Weaver *et al*, 2017). We thank Michael Hall for help with cryo-EM data collection. The electron microscopy data was collected at the Umeå Core Facility for Electron Microscopy, a node of the Cryo-EM Swedish National Facility, funded by the Knut and Alice Wallenberg, Family Erling Persson and Kempe Foundations, SciLifeLab, Stockholm University and Umeå University. This work was supported by the Deutsche Forschungsgemeinschaft (DFG) (grant WI3285/8-1 to D.N.W), the Swedish Research Council (Vetenskapsrådet) grants (2017-03783 to V.H. and 2019-01085 to G.C.A.), Ragnar Söderbergs Stiftelse (to V.H.), postdoctoral grant from the Umeå Centre for Microbial Research, UCMR (to H.T.), the European Union from the European Regional Development Fund through the Centre of Excellence in Molecular Cell Engineering (2014-2020.4.01.15-0013 to V.H.); and the Estonian Research Council (PRG335 to V.H.). D.N.W. and V.H. groups are also supported by the Deutsche Zentrum für Luft-und Raumfahrt (DLR01Kl1820 to D.N.W.) and the Swedish Research Council (2018-00956 to V.H.) within the RIBOTARGET consortium under the framework of JPIAMR.

## Materials and methods

### Strains and plasmids

All strains and plasmids used in this work are listed in Table S5.

#### E. faecalis

OG1RF and TX5332, a LsaA disruption mutant of OG1RF (Singh *et al*., 2002), were kindly provided by Dr. Barbara E. Murray (Health Science Center, University of Texas). All cloning was performed by Protein Expertise Platform at Umeå University. *E. faecalis* LsaA ORF was PCR amplified from pTEX5333 plasmid and cloned into either pCIE_cam_ (Weaver *et al*., 2017) (used for preparation of LsaA-70S complexes) or pCIE_cam_ (pCIE_cam_ derivative with the Cm^r^ gene swapped to the spectinomycin resistance Sc^r^ gene; used for MIC testing) vector for cCF10-inducible expression. To allow detection by immunoblotting and preparation of LsaA-70S complexes, the LsaA ORF was supplemented with C-terminal His_6_-TEV-FLAG_3_-tag (HTF tag) and the ribosome binding site was optimized for high expression yield. Point mutations E_142_Q and E_452_Q were introduced to LsaA resulting in pCIE_LsaA-EQ_2_-HTF.

#### S. haemolyticus

*vga(A)_LC_* gene was PCR-amplified from a *S. haemolyticus* isolate held in the O’Neill strain collection at the University of Leeds, using oligonucleotide primers vgaALC-F (5′- GGTGGT<u>GGTAC</u>CAGGATGAGGAAATATGAAAA-3′) and vgaA_LC_-R (5′- GGTGGT<u>GAATTC</u>GGTAATTTATTTATCTAAATTTCTT-3′) (engineered restriction sites shown underlined). The protein encoded by this gene is identical to that previously reported (Novotna & Janata, 2006) (accession number DQ823382). The fragment was digested with *Kpn*I and *Eco*RI and ligated into the tetracycline-inducible expression vector pRMC2 (Corrigan & Foster, 2009). Constructs encoding the VgaA_LC_ protein fused with a C-terminal FLAG_3_ tag were obtained by synthesis (Genewiz), with E_105_Q, E_410_Q and EQ_2_ mutants subsequently created by site-directed mutagenesis. Generation of other point mutants of untagged Vga(A)_LC_ was performed by NBS Biologicals, again using chemical synthesis to generate the original *vga(A)_LC_* template, followed by site-directed mutagenesis.

#### L. monocytogenes

VgaL (Lmo0919). In order to construct *L. monocytogenes* EDGe::Δ*lmo0919*, regions corresponding to the upstream and downstream flanking regions of *lmo0919*, present on the EDGe genome were amplified with primer pairs VKT35 (5′- GGGGGGATCCATCACTAGCCGAATCCAAAC-3′), VKT36 (5′- GGGGGAATTCAAAAAATAACCTCCTGAATATTTTCAGAG-3′) and VHKT37 (5′- GGGGGAATTCATTGTTGTCTTTTTATTCAAGCTAAATAAAAAA-3′), VHKT38 (5′- GGGGCCATGGCGTGCTGTACGGTATGC-3′) respectively. Fragments were then cloned in tandem into the pMAD vector using *Bam*HI, *Eco*RI and *Nco*RI restriction sites. The resulting vector, VHp689, was then sequenced to ensure wild-type sequences of clones. Gene deletion was then performed as per Arnaud *et. al.* (Arnaud *et al*, 2004).

*lmo0919* was amplified from EDGe genomic DNA using primers VHKT12 (5′- CCCCCCATGGCATCTACAATCGAAATAAATC-3′) and VHKT39 (5′- GGGGCTGCAGTTAACTAAATTGCTGTCTTTTTG-3′), and cloned into pIMK3 using *Nco*I and *Pst*I restriction sites, resulting in plasmid VHp690.

Overlap extension PCR was used in order to introduce a HTF tag at the C-terminus of *lmo0919* (Ho *et al*, 1989). The *lmo0919* locus and HTF tag were amplified with primer pairs VHKT12, VHKT15 (5′-ATGATGATGGCCGCCACTAAATTGCTGTCTTTTTG-3′) and VHKT14 (5′- AGACAGCAATTTAGTGGCGGCCATCATCATCATC-3′), VHKT13 (5′- GGGGCTGCAGTTAGCCTTTGTCATCGTC-3′) using EDGe genomic DNA and VHp100 template DNA respectively, producing fragments with overlapping ends. VHKT12 and VHKT13 were then used to fuse the fragments and the resulting PCR product was cloned into pIMK3 using *Nco*I and *Pst*I sites resulting in VHp692.

To introduce EQ2 mutations (E104Q and E408Q) simultaneously into the VHp692 plasmid, primers VHT266 (5′-TCTTGATCAACCAACCAACTATTTGGATATCTACGCAATGGAA-3′) and VHT267 (5′-TTGTTGGTTGGTCTGCTAGGAGAACACTTGGATTTTGGCGCA-3′) containing both mutations were used to extend out from *lmo0919^HTF^* to amplify the VHp692 backbone. Primers VHT264 (5′-AGCAGACCAACCAACAAGCAATCTTGATGTCG-3′) and VHT265 (5′-TGGTTGGTTGATCAAGAATCAAGAAATTGGCGT-3′) also containing *lmo0919^EQ2^* mutations were used to amplify a fragment with overlapping sequence to the backbone fragment. Both PCR products were then assembled using NEBuilder® HiFi DNA Assembly Master Mix (NEB), resulting in VHp693.

#### B. subtilis

To construct the VHB109 [*trpC2* Δ*vmlR* thrC::P*_hy-spnak_*-*lsaA kmR*] strain untagged LsaA under the control of an IPTG-inducible P*_hy-spank_* promotor, a PCR product encoding lsa(A) was PCR-amplified from pTEX5333 using the primers VHT127 (5′- CGACGAAGGAGAGAGCGATAATGTCGAAAATTGAACTAAAACAACTATC-3′) and VHT128 (5′-CACCGAATTAGCTTGCATGCTTATGATTTCAAGACAATTTTTTTATCTGTTA-3′). The second PCR fragment encoding a kanamycin-resistance marker, a polylinker downstream of the Phy-spank promoter and the lac repressor ORF – all inserted in the middle of the thrC gene – was PCR-amplified from pHT009 plasmid using primers VHT123 (5′- CATTATCGCTCTCTCCTTCGTCGACTAAGCTAATTG-3′) and VHT125 (5′-TAAGCATGCAAGCTAATTCGGTGGAAACGAGG-3′). The two fragments were ligated using the NEBuilder HiFi DNA Assembly master mix (New England BioLabs, Ipswich, MA) yielding the pHT009-lsaA plasmid (VHp369) which was used to transform the VHB5 [*trpC2* Δ*vmlR*] strain. Selection for kanamycin resistance yielded the desired VHB109 strain. To construct the VHB168 [*trpC2* Δ*vmlR* thrC::P*_hy-spnak_-lsaAK244A kmR*] strain, VHp369 plasmid was subjected to site-directed mutagenesis using primer VHP303 (5′- GCATCACCTTCACGGTTCATCGACCATTCCGCT-3′) and VHP304 (5′-GTACGGCAACGCTAAGGAAAAAGGGAGCGGGGCGA-3′), according to directions of Phusion Site-Directed Mutagenesis Kit (Thermo Fisher Scientific), yielding, yielding VHp526 (pHT009-*lsaAK244A*) plasmid which was used to transform the VHB5 [*trpC2* Δ*vmlR*] strain. Selection for kanamycin resistance yielded the desired VHB168 strain. To construct the VHB169 [*trpC2* Δ*vmlR thrC*::P*_hy-spnak_*-*lsaAF257A kmR*] strain, VHp369 plasmid was subjected to site-directed mutagenesis using primer VHP305 (5′- CAATCGCCCCGCTCCCTTTTTCCTTAGCGT-3′) and VHP306 (5′-CGGATACAGGAGCCATTGGTGCCCGGGCA-3′), according to directions of Phusion Site-Directed Mutagenesis Kit (Thermo Fisher Scientific), yielding, yielding VHp527 (pHT009-*lsaAF257A*) plasmid which was used to transform the VHB5 [*trpC2* Δ*vmlR*] strain. Selection for kanamycin resistance yielded the desired VHB169 strain.

### Bacterial transformation

#### E. faecalis

Electrocompetent cells were prepared as per Bhardwaj and colleagues (Bhardwaj *et al*, 2016). Shortly, an over-night culture grown in the presence of appropriate antibiotics was diluted to OD_600_ of 0.05 in 50 mL of BHI media (supplemented with 2 mg/mL kanamycin in case of TX5332), grown to OD_600_ of 0.6-0.7 at 37 °C with moderate shaking (160 rpm). Cells were collected by centrifugation at 4,000 rpm at 4 °C for 10 min. Cells were resuspended in 0.5 mL of sterile lysozyme buffer (10 mM Tris-HCl pH 8; 50 mM NaCl, 10 mM EDTA, 35 µg/mL lysozyme), transferred to 1.5 mL Eppendorf tube and incubated at 37 °C for 30 minutes. Cells were pelleted at 10,000 rpm at 4 °C for 10 min and washed three times with 1.5 mL of ice-cold electroporation buffer (0.5M sucrose, 10% glycerol(w/v)). After last wash the cells were resuspended in 500 µL of ice-cold electroporation buffer and aliquoted and stored at –80°C. For electroporation 35 µL of electrocompetent cells were supplemented with 1 µg of plasmid DNA, transferred to ice-cold 1 mm electroporation cuvette and electroporated at 1.8 keV. Immediately after electroporation 1 mL of ice-cold BHI was added to the cells, the content of the cuvette was transferred to 1.5 mL Eppendorf tubes and the cells were recovered at 37 °C for 2.5 hours and plated to BHI plates containing appropriate antibiotics (10 µg/mL chloramphenicol and 2 mg/mL kanamycin).

#### S. aureus

the preparation and transformation of *S. aureus* electrocompetent cells followed the method of Schenk & Laddaga (Schenk & Laddaga, 1992), though used Tryptone soya broth (Oxoid) containing 2.5% yeast extract in place of B2 medium. Sequence-verified constructs established in *E. coli* were transferred into the restriction deficient *S. aureus* RN4220 strain (Fairweather *et al*, 1983), before recovery and introduction into *S. aureus* SH1000 (Horsburgh *et al*, 2002; O’Neill, 2010).

#### L. monocytogenes

pIMK3 integrative plasmids were transformed into *L. monocytogenes* via conjugation. *E. coli* S17.1 harbouring pIMK3 and its derivatives, was grown at 37 °C overnight in LB media supplemented with 50 µg/mL Kanamycin, 1 mL of culture was washed three times with sterile BHI media to remove antibiotics. 200 µL of washed *E. coli* culture was mixed with an equal volume of *L. monocytogenes* overnight culture grown at 37 °C in BHI media. 200 µl of mixed bacterial suspension was then dropped onto a conjugation filter (Millipore #HAEP047S0) placed onto a BHI agar plate containing 0.2 µg/mL penicillin-G. After overnight incubation at 37 °C, bacterial growth from the filter was re-suspended in 1 ml of BHI and 100-300 µL plated onto BHI-agar plates supplemented with 50 µg/mL Kanamycin (to select for pIMK3), 50 µg/mL Nalidixic acid and 10 µg/mL Colistin sulfate (Sigma-Aldrich C4461-100MG). Resulting colonies were checked for correct integration via PCR and subsequent sequencing using primers VHKT42 and VHKT43.

### Antibiotic susceptibility testing

Minimum Inhibitory Concentrations (MIC) were determined based on guidelines from the European Committee on Antimicrobial Susceptibility Testing (EUCAST) (http://www.eucast.org/ast_of_bacteria/mic_determination).

#### E. faecalis

bacteria were grown in BHI media supplemented with 2 mg/mL kanamycin (to prevent *lsa* revertants), either 0.1 mg/mL spectinomycin (to maintain the pCIE_spec_ plasmid),) or 20 µg/mL of chloramphenicol (to maintain the pCIE_cam_ plasmid used to validate the functionality of the HTF-tagged LsaA variant), 100 ng/mL of cCF10 peptide (to induce expression of LsaA protein) as well as increasing concentrations of antibiotics was inoculated with 5 × 10^5^ CFU/mL (OD_600_ of approximately 0.0005) of *E. faecalis* Δ*lsaA* (*lsa::Kan*) strain TX5332 transformed either with empty pCIE_spec_ plasmid, or with pCIEspec encoding LsaA. After 16-20 hours at 37 °C without shaking, the presence or absence of bacterial growth was scored by eye.

#### S. aureus

bacteria were grown in cation-adjusted Mueller-Hinton Broth (MHB) at 37 °C with vigorous aeration, supplemented with 10 mg/L chloramphenicol to maintain the pRMC2 plasmid. Upon reaching an absorbance of OD_625_ of 0.6, anhydrotetracycline (ATC) (Sigma-Aldrich, UK) was added at a final concentration of 100 ng/mL to induce expression from pRMC2, and incubated for a further 3 hours. Cultures were then diluted to 5 x 10^5^ CFU/mL using MHB supplemented with ATC (100 ng/mL) and used in MIC determinations essentially as described above (though cultures were shaken).

#### L. monocytogenes

bacteria were grown in BHI media supplemented with 50 µg/mL kanamycin (to prevent loss of the integrated pIMK3 plasmid), 1 mM of IPTG (to induce expression of VgaL protein) as well as increasing concentrations of antibiotics was inoculated with 5 x 10^5^ CFU/mL (OD_600_ of approximately 0.0003) of *L. monocytogenes* EDG*-e* wildtype strain or EDG-*e*::Δ*lmo0919* strain transformed either with empty pIMK3 plasmid, or with pIMK3 encoding VgaL variants. After 16–20 hours at 37 °C without shaking, the presence or absence of bacterial growth was scored by eye.

#### B. subtilis

(for LsaA mutants): *B. subtilis* strains were pre-grown on LB plates either supplemented with 1 mM IPTG overnight at 30 °C. Fresh individual colonies were used to inoculate filtered LB medium in the presence of 1 mM IPTG, and OD_600_ adjusted to 0.01. The cultures were seeded on a 100-well honeycomb plate (Oy Growth Curves AB Ltd, Helsinki, Finland), and plates incubated in a Bioscreen C (Labsystems, Helsinki, Finland) at 37°C with continuous medium shaking. After 90 min (OD_600_ ≍ 0.1), antibiotics were added and growth was followed for an additional 6 hours.

### Preparation of bacterial lysates

#### Preparation of bacterial biomass

##### E. faecalis

*E. faecalis* TX5332 transformed with pCIE plasmids (either empty vector and expressing either wild-type or EQ_2_ variants of C-terminally HTF-tagged LsaA) were grown overnight from single colony in BHI supplemented with 2 mg/mL kanamycin and 10 µg/mL of chloramphenicol. Next day overnight cultures were diluted to starting OD_600_ of 0.05 in 160 mL BHI supplemented with 0.5 mg/mL kanamycin and 10 µg/mL of chloramphenicol. Cells were grown with intensive shaking at 37 °C till OD_600_ of 0.6 and were induced with 300 ng/mL of cCF10 peptide for 30 minutes prior harvesting by centrifugation at 10,000 ×g for 15 minutes at 4 °C.

##### S. aureus

*S. aureus* SH1000 transformed with pRMC2 plasmids (empty vector, wild-type and EQ_2_ VgaA_LC_-FLAG_3_) were grown in LB supplemented with 25 µg/mL of chloramphenicol. Saturated cultures were diluted to an OD_600_ of 0.1 in 400 mL LB supplemented with 20 µg/mL of chloramphenicol and grown at 37 °C with vigorous aeration to an OD_600_ of 0.6. Protein expression was induced with 100 ng/mL of anhydro-tetracycline for 30 minutes prior to harvesting by centrifugation at 10 000 ×g for 15 minutes at 4 °C.

##### L. monocytogenes

*L. monocytogenes* EDG-*e* was transformed with pIMK3 plasmids (empty vector, wild-type and EQ_2_ VgaL-HTF) were grown overnight from single colony in LB supplemented with 50 µg/mL of kanamycin. Next day overnight cultures were diluted till starting OD_600_ of 0.005 in 200 mL BHI supplemented with 50 µg/mL of Kanamycin. Cells were grown at 37 °C with shaking at 160 rpm till OD_600_ of 0.6 and were induced with 1 mM IPTG for 60 minutes prior harvesting by centrifugation at 10,000 ×g for 15 minutes at 4 °C.

#### Preparation of clarified lysates

Cell pellets were resuspended in 1.5 mL of cell lysis buffer (95 mM KCl, 5 mM NH_4_Cl, 20 mM HEPES pH 7.5, 1 mM DTT, 5 mM Mg(OAc)_2_, 0.5 mM CaCl_2_, 8 mM putrescine, 1 mM spermidine, 1 tablet of cOmplete™ EDTA-free Protease Inhibitor Cocktail (Roche) per 10 mL of buffer and in the absence or presence of either 0.5 or 0.75 mM ATP), resuspended cells were opened by FastPrep homogeniser (MP Biomedicals) with 0.1 mm Zirconium beads (Techtum) in 4 cycles by 20 seconds with 1 minute chill on ice. Cell debris was removed after centrifugation at 14,800 ×g for 15 minutes at 4 °C. Total protein concentration in supernatant was measured by Bradford assay (BioRad), supernatant was aliquoted and frozen in liquid nitrogen.

### Polysome fractionation and immunoblotting

#### Sucrose density gradient centrifugation

After melting the frozen lysates on ice, 2 A_260_ units of each extract was aliquoted into three tubes and supplemented with or without 0.5-0.75 mM ATP and was loaded onto 5–25% or 7– 35% (w/v) sucrose density gradients in HEPES:Polymix buffer (Takada *et al*, 2020), 5 mM Mg(OAc)_2_ and supplemented or not with 0.5–0.75 mM ATP. Gradients were resolved at 35,000 rpm for 2.5 hours at 4 °C in SW41 rotor (Beckman) and analysed and fractionated using Biocomp Gradient Station (BioComp Instruments) with A_280_ as a readout.

#### Immunoblotting

##### LsaA and VgaA_LC_

Schleicher & Schuell Minifold II Slot Blot System SRC072/0 44-27570 Manifold was used for transferring samples from sucrose gradient fractions to PVDF membranes (Immobilon PSQ, Merk Millipore). Shortly, 15-100 μL of each sucrose gradient fraction was added to 200 μL of Slot-blotting Buffer (20 mM HEPES:KOH pH 7.5, 95 mM KCl, 5 mM NH_4_Cl, 5 mM Mg(OAc)_2_) in slots and blotted onto PVDF membrane that had been activated with methanol for one minute, wetted in MilliQ water and equilibrated with Slot-blotting Buffer (1x PM 5 mM Mg^2+^ without putrescine and spermidine) for 10 minutes. After blotting of the samples each slot was washed twice with 200 μL of Slot-blotting Buffer. The membrane was removed from the blotter, transferred to hybridization bottle, equilibrated for 10 minutes in PBS-T (1x PBS supplemented with 0.05% Tween-20) and blocked in PBS-T supplemented with 5% w/v nonfat dry milk for one hour. Antibody incubations were performed for one hour in 1% nonfat dry milk in PBS-T with five 5-minute washes in fresh PBS-T between and after antibody incubations. HTF-tagged LsaA and FLAG_3_-tagged VgaA_LC_ proteins were detected using anti-Flag M2 primary (Sigma-Aldrich, F1804; 1:10,000 dilution) antibodies combined with anti-mouse-HRP secondary (Rockland; 610-103-040; 1:10,000 dilution) antibodies. An ECL detection was performed on ImageQuant LAS 4000 (GE Healthcare) imaging system using Pierce® ECL Western blotting substrate (Thermo Scientific). The blotting and all incubations were performed at room temperature in hybridization oven.

##### VgaL (Lmo0919)

Western blotting of lysates on sucrose gradient fractionation was performed as previously described (Takada *et al*., 2020). VgaL-HTF was detected using anti-Flag M2 primary (Sigma-Aldrich, F1804; 1:10,000 dilution) antibodies combined with anti-mouse-HRP secondary (Rockland; 610-103-040; 1:10,000 dilution) antibodies.

### Affinity purification on anti-FLAG M2 affinity gel

100 µL of well mixed anti-FLAG M2 Affinity Gel aliquots were loaded on columns (Micro Bio-Spin Columns, Bio-Rad) and washed two times with 1 mL of cell lysis buffer by gravity flow. All incubations, washings and elutions were done at 4 °C.

The total protein concentration of each lysate was adjusted to 2 mg/mL with cell lysis buffer and 1 mL of each lysate was loaded on columns and incubated for two hours with end-over-end mixing for binding. The columns were washed 5 times by 1 mL of cell lysis buffer by gravity flow. For elution of FLAG-tagged proteins and their complexes 100-300 µL of 0.1 mg/mL FLAG_3_ peptide (Sigma) was added to samples, the solutions were incubated at 4 °C for 20 minutes with end-over-end mixing. Elutions were collected by centrifugation at 2,000 ×g for 2 minutes at 4 °C.

20 µL aliquots of collected samples (flow-through, washes and elutions) were mixed with 5 µL of 5x SDS loading buffer and heated up at 95 °C for 15 minutes. The beads remaining in the column were washed twice with 1 mL of cell lysis buffer and resuspended in 100 µL of 1x SDS loading buffer. Denatured samples were resolved on 12-15% SDS-PAGE. SDS-gels were stained by “Blue-Silver” Coomassie Staining (Candiano *et al*, 2004) and washed with water for 6 hours or overnight before imaging with LAS4000 (GE Healthcare).

### tRNA microarrays

To fully deacylate tRNAs, eluates and input lysate samples from two biological replicates were mixed with 80 µL 250 mM Tris-HCl, pH 9.0, 10 µL 0.2 M EDTA, 10 µL 1% SDS, and incubated for 45 min, and neutralised with 200 µL 1 M NaOAc, pH 5.5, before mixing 1:1 with acidic phenol:chloroform alcohol 5:1. The supernatant was precipitated with ethanol and dissolved in ddH_2_O.

tRNA microarrays were performed as described (Kirchner *et al*, 2017). Briefly, using the unique invariant single stranded 3′-NCCA-ends of intact tRNA a Cy3-labelled or Atto647-labeled RNA/DNA hybrid oligonucleotide was ligated to the tRNA extracted from the RqcH-50S samples and total *E. faecalis* tRNA (from the lysate), respectively. Labeled tRNA was purified by phenol:chloroform extraction and loaded on a microarray containing 24 replicates of full-length tDNA probes recognizing *E. faecalis* tRNA isoacceptors. Florescence signals were normalized to three *in vitro* transcribed human tRNAs, spiked in to each sample. Microarrays were statistically analysed with in-house scripts written in Python 3.7.0.

### Grid preparation, cryo-electron microscopy and single-particle reconstruction

#### Preparation of cryo-EM grids and data collection

Elutions from LsaA and VgaL pull-downs were loaded on grids within two hours after obtaining them without freezing, samples were kept on ice. The VgaA_LC_ sample was frozen in liquid nitrogen after pull-down, defrosted and loaded later. After glow-discharging of grids, 3.5 mL of sample was loaded on grids in Vitrobot (FEI) in conditions of 100% humidity at 4 °C, blotted for 5 seconds and vitrified by plunge-freezing in liquid ethane. Samples were imaged on a Titan Krios (FEI) operated at 300 kV at a nominal magnification of 165 k× (0.86 Å/pixel, later estimated to be 0.82 Å/pixel by comparing refined maps to structures with known magnification) with a Gatan K2 Summit camera at an exposure rate of 5.80 electrons/pixel/s with a 4 seconds exposure and 20 frames using the EPU software. Quantifoil 1.2/1.3 Cu_200_ grids were used for LsaA and VgaA_LC_ and Quantifoil 2/2 Cu_200_ grids were used for VgaL.

#### Single-particle reconstruction

Motion correction was performed with MotionCor2 with 5×5 patches (Zheng *et al*, 2017). Relion 3.0 or 3.1 was used for further processing unless otherwise stated and resolutions are reported according to the so-called ‘gold standard’ criteria (Henderson *et al*, 2012; Scheres & Chen, 2012; Zivanov *et al*, 2018). CTFFIND4 (LsaA dataset) or Gctf v1.06 (VgaA_LC_ and VgaL datasets) was used for CTF estimation (Rohou & Grigorieff, 2015; Zhang, 2016). Particles were picked with Gautomatch (https://www2.mrc-lmb.cam.ac.uk/research/locally-developed-software/zhang-software/#gauto, developed by K. Zhang) without supplying a reference, and in the case of LsaA, re-picked using RELION autopicker after templates were generated by 2D classification. Particles were initially extracted at 2.46 Å/pixel and subjected to 2D classification. Classes that resembled ribosomes were used for 3D refinement, with a 60 Å low-pass filter applied to initial references. For 3D refinement of LsaA–70S, the initial reference was EMDB-0176, a *B. subtilis* 70S ribosome with no factor bound in the E-site (Crowe-McAuliffe *et al*., 2018); for VgaA_LC_–70S and VgaL–70S 3D refinements the RELION initial model job type was used to create a reference from particles selected after 2D classification. 3D classification was performed without angular sampling, and classes of interest were re-extracted at 0.82 Å/pixel for further refinement.

In the case of LsaA, after initial 3D classification, a soft mask around the A-site was used for partial signal subtraction followed by focussed classification. The classes with the strongest and weakest A-site density were selected for signal restoration and refinement. In the case of the VgaA_LC_ dataset, initial 3D classification yielded a class with apparent sub-stoichiometric density in the E-site corresponding to VgaA_LC_. Micrographs with poor values from CTF estimation were discarded, particles were re-extracted, subjected to an additional 2D classification and 3D refinement, followed by Bayesian polishing and CTF refinement. An additional 3D classification yielded a class with strong E-site density corresponding to the factor. Refer to Figures S4–S6 for details.

For multibody refinements, soft masks around the small subunit body, small subunit head, and large subunit/ARD were applied. In the case of the VgaA_LC_ dataset, particles were first re-extracted in a smaller box (360×360 pixels) and subjected to 3D refinement prior to multibody refinement.

ResMap was used to estimate local resolution (Kucukelbir *et al*, 2014). Maps were locally filtered using SPHIRE (Moriya *et al*, 2017).

#### Molecular modelling

For the *E. faecalis* and *L. monocytogenes* ribosomes, homology models were generated with SWISS-MODEL (Waterhouse *et al*, 2018), mostly from PDB 6HA1/6HA8 (Crowe-McAuliffe *et al*., 2018). PDBs 4YBB (Noeske *et al*, 2015) 5MDV (James *et al*, 2016) were used as additional templates and references where necessary, 4V9O (Pulk & Cate, 2013) was used for bS21, 5ML7 (Gabdulkhakov *et al*, 2017) and 3U4M (Tishchenko *et al*, 2012) were used for the L1 stalk region, 5AFI (Fischer *et al*, 2015) and 5UYQ (Loveland *et al*, 2017) were used for tRNAs, and 6QNQ was used to help placing metal ions (Rozov *et al*, 2019). PDB 5LI0 (Khusainov *et al*., 2016) was used as a starting model for the *S. aureus* ribosome. Where appropriate, individual components of multibody refinements were fitted into density from the corresponding locally filtered map to help modelling. Models were adjusted with Coot (Casañal *et al*, 2020) and refined using locally filtered maps in Phenix version 1.14 3260 (Liebschner *et al*, 2019).

Figures were created with PyMOL 2.0 (Schrödinger, LLC), UCSF Chimera (Pettersen *et al*, 2004), RELION (Zivanov *et al*., 2018), and Igor Pro (WaveMetrics, Inc.). Structures were aligned in PyMOL using the 23S rRNA unless otherwise noted.

Figures were assembled with Adobe Illustrator (Adobe Inc.).

## Supplementary Material

**Figure S1.**
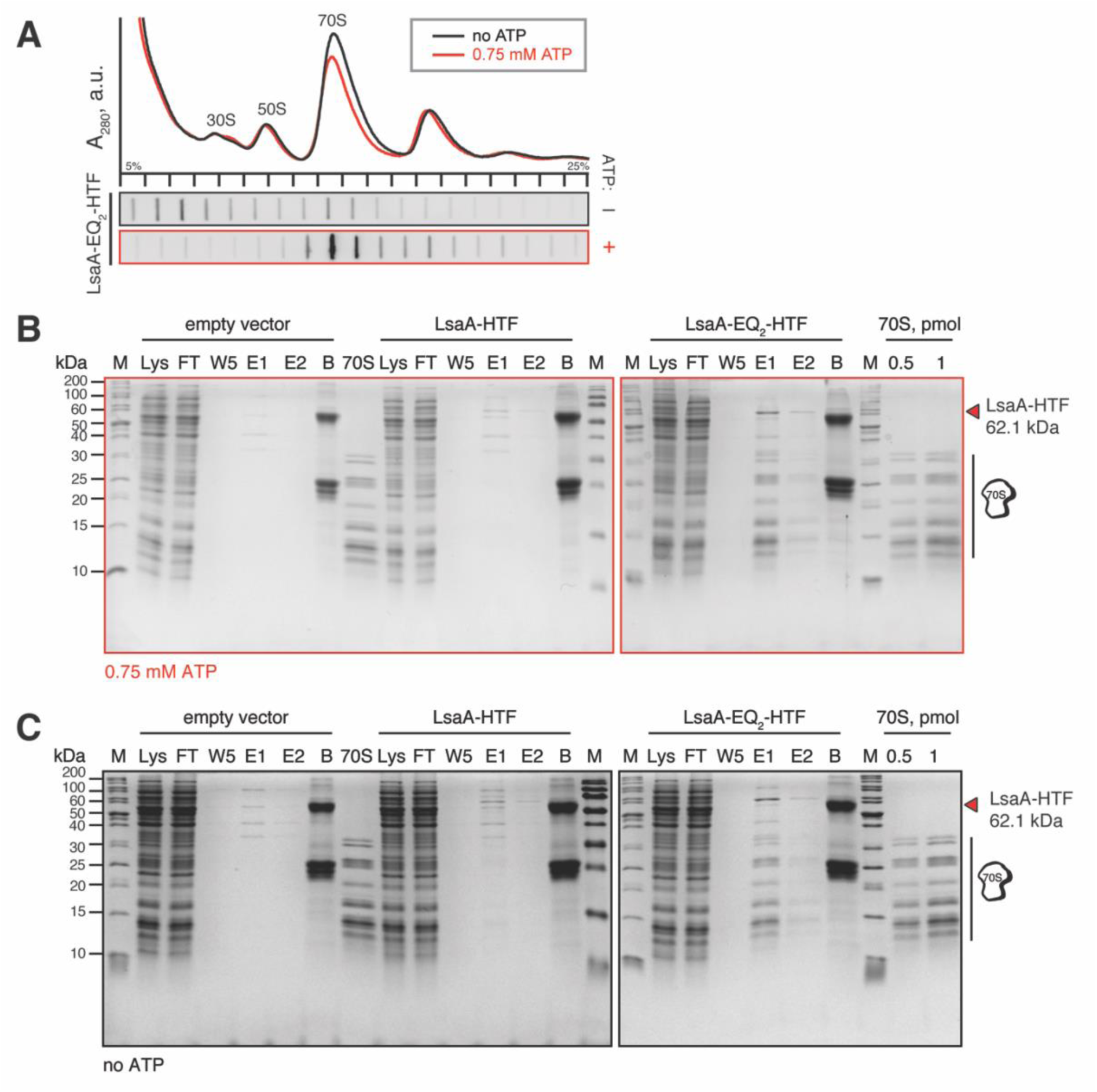
Characterization of *E. faecalis* LsaA interactions with ribosomes and preparation of samples for cryo-EM. (**A**) Polysome profiles and immunoblot analyses of C-terminally His_6_-TEV-FLAG_3_-tagged (HTF) ATPase-deficient (EQ_2_) LsaA-EQ_2_ ectopically expressed in Δ*lsaA E. faecalis* TX5332. Experiments were performed both in the presence or absence of 0.75 mM ATP in gradients. (**B, C**) Affinity purification of wild-type and EQ_2_ *E. faecalis* LsaA-HTF ectopically expressed in TX5332 *E. faecalis*. Pull-down experiments were performed either in the presence (**B**) or absence (**C**) of 0.75 mM ATP using clarified lysates of *E. faecalis* either transformed with empty pCIE vector (background control), expressing *E. faecalis* LsaA-HTF (VHp100) or expressing *E. faecalis* LsaA-EQ_2_-HTF (VHp149). Samples: M: molecular weight marker; Lys: 2 μL of clarified lysate, FT: 2 μL of flow-through; W5: 10 μL of last wash before specific elution; E1: 10 μL of the first elution with FLAG_3_ peptide; E2: 10 μL of the second elution with FLAG_3_ peptide; B: 10 μL of SDS-treated post-elution anti-FLAG beads; 70S: purified *E. faecalis* 70S ribosomes. The samples were resolved on 15% SDS-PAGE gel. The 0.75 mM ATP *E. faecalis* LsaA-EQ_2_-HTF pulldown sample was used for further cryo-EM and tRNA array analysis.

**Figure S2.**
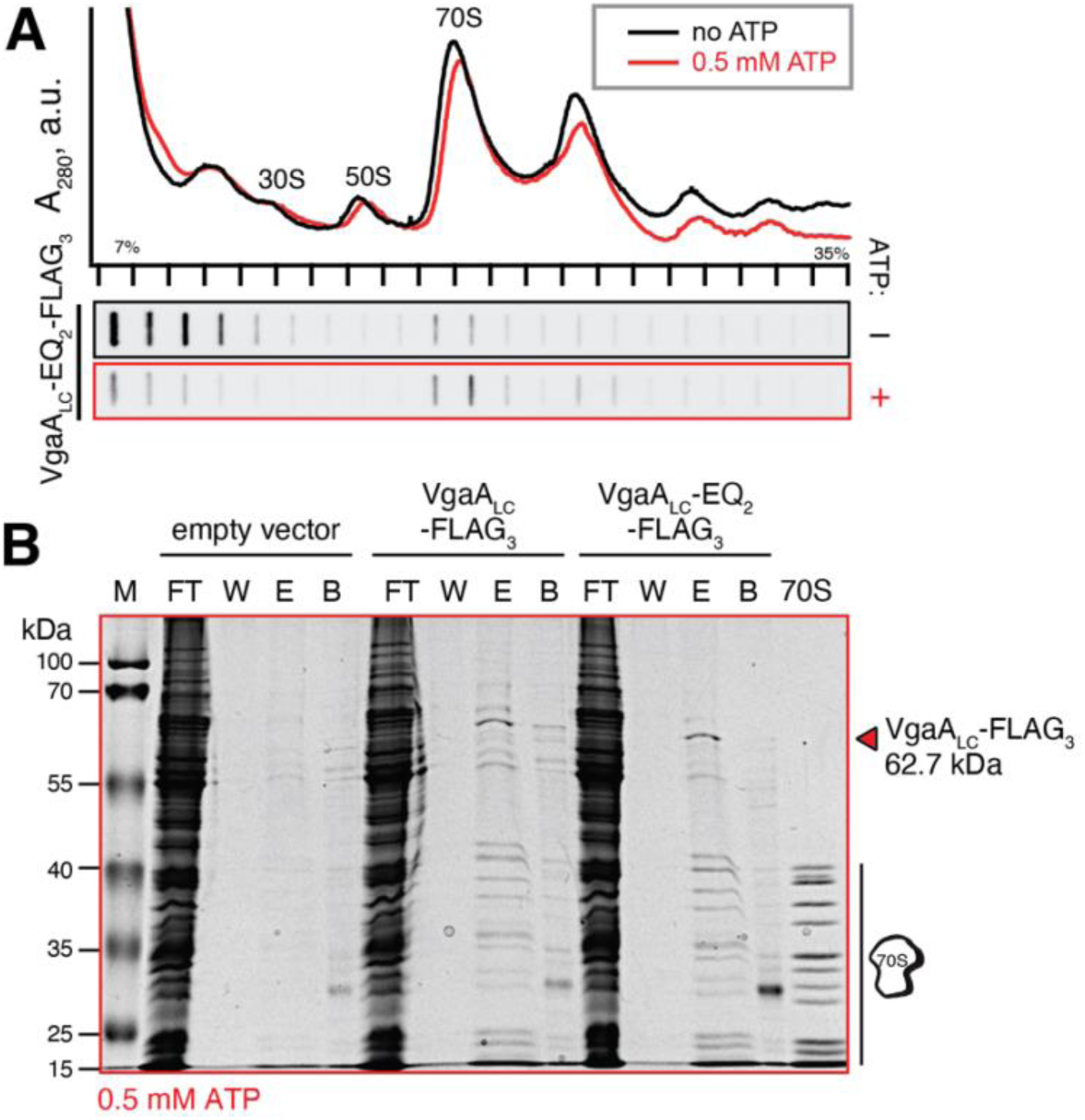
Characterization of *S. haemolyticus* VgaA_LC_ interactions with ribosomes and reparation of samples for cryo-EM reconstructions. **(A)** Polysome profiles and immunoblot analyses of FLAG_3_-tagged *S. haemolytic_us_* VgaA_LC_-EQ_2_ ectopically expressed in wild-type SH-1000 *S. aureus*. Experiments were performed both in the presence or absence of 0.5 mM ATP in gradients. **(B)** Affinity purification of wild-type and EQ_2_ *S._h_ aemolyticus* VgaA_LC_-FLAG_3_ ectopically expressed in SH-1000 *S. aureus*. Immunoprecipitations were performed in the presence of 0.5 mM ATP and the samples were resolved on a 15% polyacrylamide gel by SDS-PAGE. Samples: M: 2 μL of molecular weight marker; FT: 2 μL of flow-through, W: 10 μL of last wash before specific elution; E: 10 μL of elution with FLAG_3_ peptide; B: 2 μL of SDS-treated post-elution anti-FLAG beads; 70S: 1 pmol of purified *S. aureus* 70S ribosomes. The 0.5 m_M_ ATP *S. haemolyticus* VgaA_LC_-EQ_2_-HTF pulldown sample was used for cryo-EM reconstructions.

**Figure S3.**
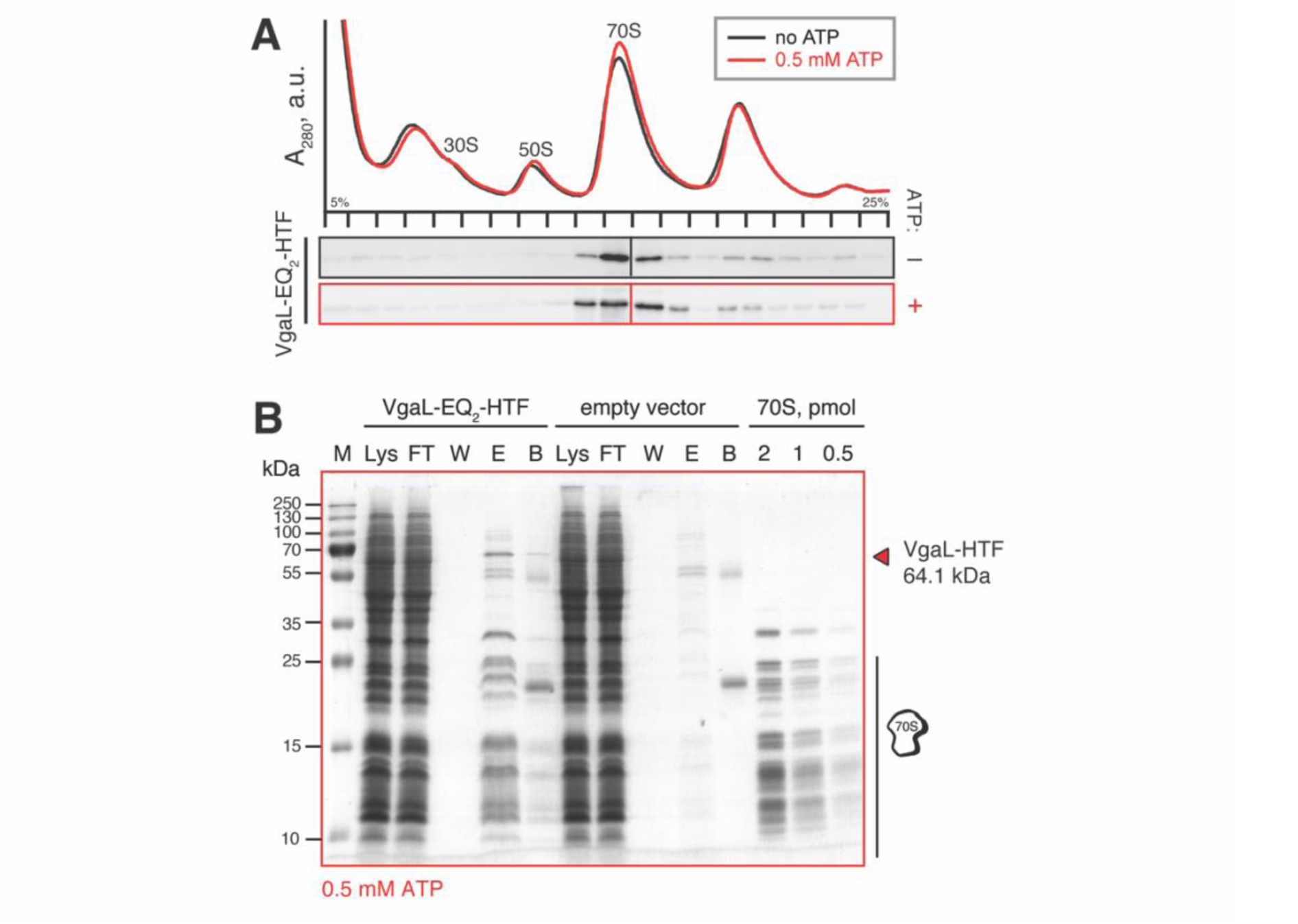
Characterization of *L. monocytogenes* VgaL (Lmo0919) interactions with ribosomes and reparation of samples for cryo-EM reconstructions. **(A)** Polysome profiles and immunoblot analyses of HTF-tagged *L. monocytogenes* VgaL-EQ_2_ (Lmo0919-EQ_2_) ectopically expressed in wild-type EDG-*e L. monocytogenes*. Experiments were performed both in the presence or absence of 0.5 mM ATP in gradients. **(B)** Affinity purification of *L. monocytogenes* VgaL-EQ_2_ ectopically expressed in EDG-*e L. monocytogenes*. Pull-down experiments were performed in the presence of 0.5 mM ATP using clarified lysates of *L. monocytogenes* transformed with empty integrative pIMK3 vector (background control), expressing VgaL-HTF (VHp692) or expressing VgaL-EQ_2_-HTF (VHp149). Samples: M: 2 μL of molecular weight marker; FT: 2 μL of flow-through; W: 10 μL of last wash before specific elution; E: 10 μL of elution with FLAG_3_ peptide; B: 2 μL of SDS-treated post-elution anti-FLAG beads; 70S: purified *B*. *subtilis* 70S ribosomes, the samples were resolved on 15 % SDS-PAGE gel. The 0.5 mM ATP *L. monocytogenes* VgaL-EQ_2_-HTF pulldown sample was used for cryo-EM reconstructions.

**Figure S4.**
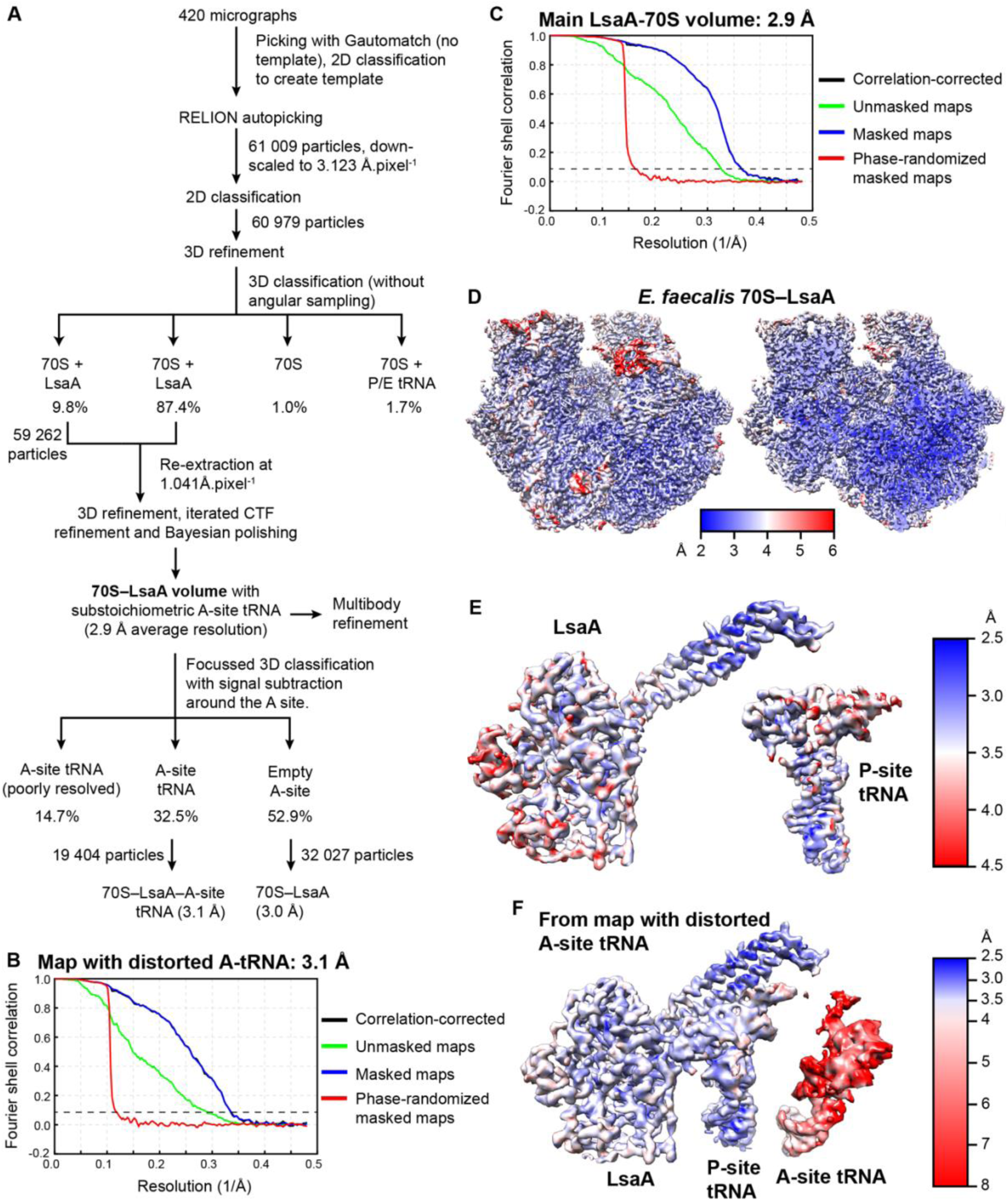
Processing of the cryo-EM data of LsaA-70S complex. **(A)** Processing scheme for the LsaA-70S complex, yielding two subpopulations of LsaA-70S complexes with and without A-site tRNA. **(B, C)** Fourier Shell Correlation (FSC) curves of the LsaA-70S **(B)** with A-tRNA and **(C)** without A-tRNA with a dashed line at 0.143 indicating average resolutions of 3.1 Å and 2.9 Å, respectively. **(D)** Overview (left) and transverse section (right) of the cryo-EM map of the LsaA-70S (without A-tRNA) coloured according to local resolution. **(E)** Isolated density of LsaA (left) and P-site tRNA (right) from **(D)**. **(F)** Isolated density of LsaA, P-site and A-site tRNA from the LsaA-70S map (with A-tRNA) coloured according to local resolution.

**Figure S5.**
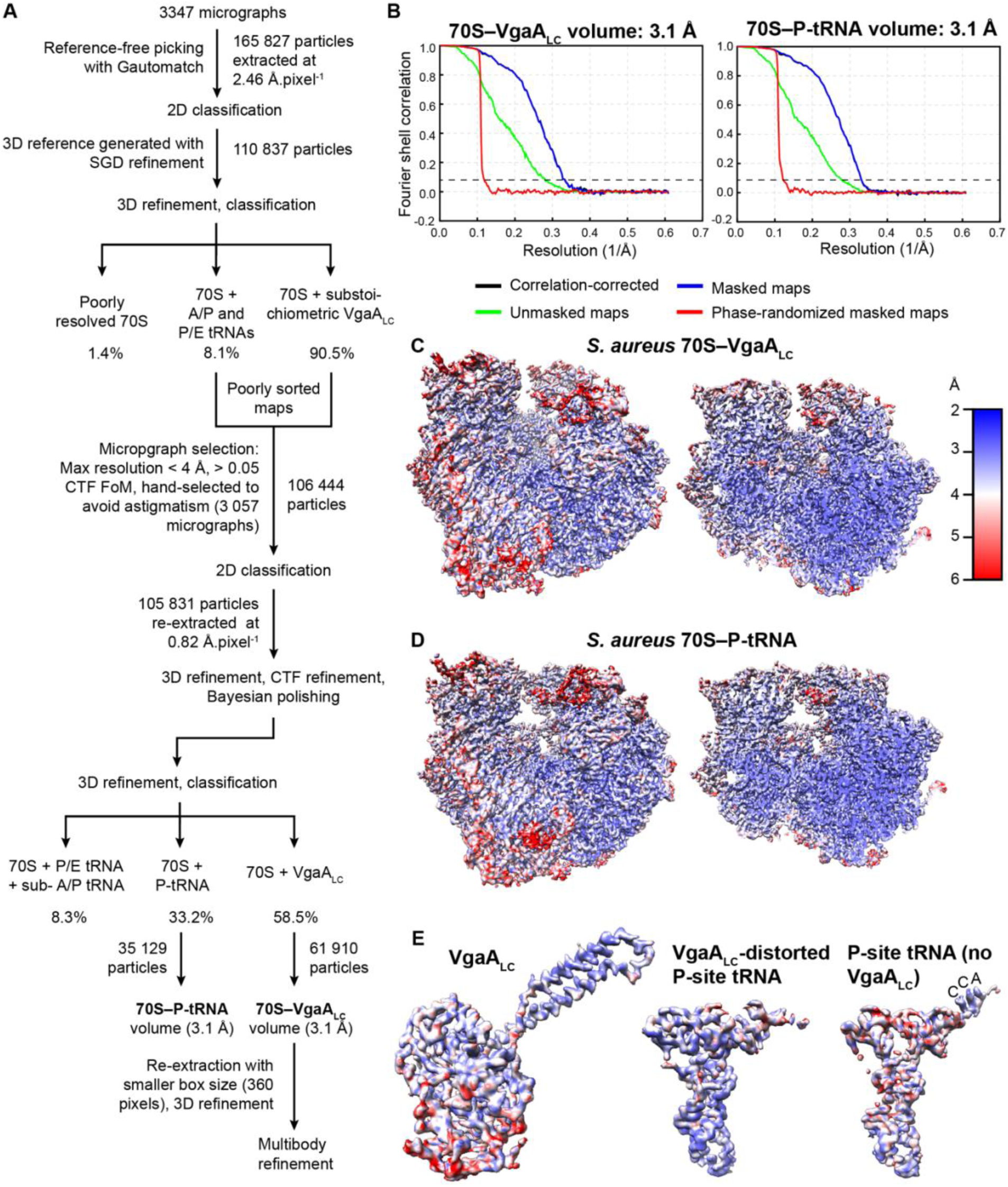
Processing of the cryo-EM data of VgaA_LC_–70S complex. **(A)** Processing scheme for the LsaA-70S complex, yielding a VgaA_LC_–70S and 70S–P-tRNA complex without VgaA_LC_. **(B)** Fourier Shell Correlation (FSC) curves of the VgaA_LC_–70S and 70S–P-tRNA complexes with a dashed line at 0.143 indicating average resolutions of 3.1 Å. **(C, D)** Overview (left) and transverse section (right) of the cryo-EM map of the **(C)** VgaA_LC_–70S and **(D)** 70S-P-tRNA complexes coloured according to local resolution. **(E)** Isolated density of VgaA_LC_ (left) and P-site tRNA (right) from the VgaA_LC_–70S complex, and the P-site-tRNA from the 70S–P-tRNA complex, coloured according to local resolution.

**Figure S6.**
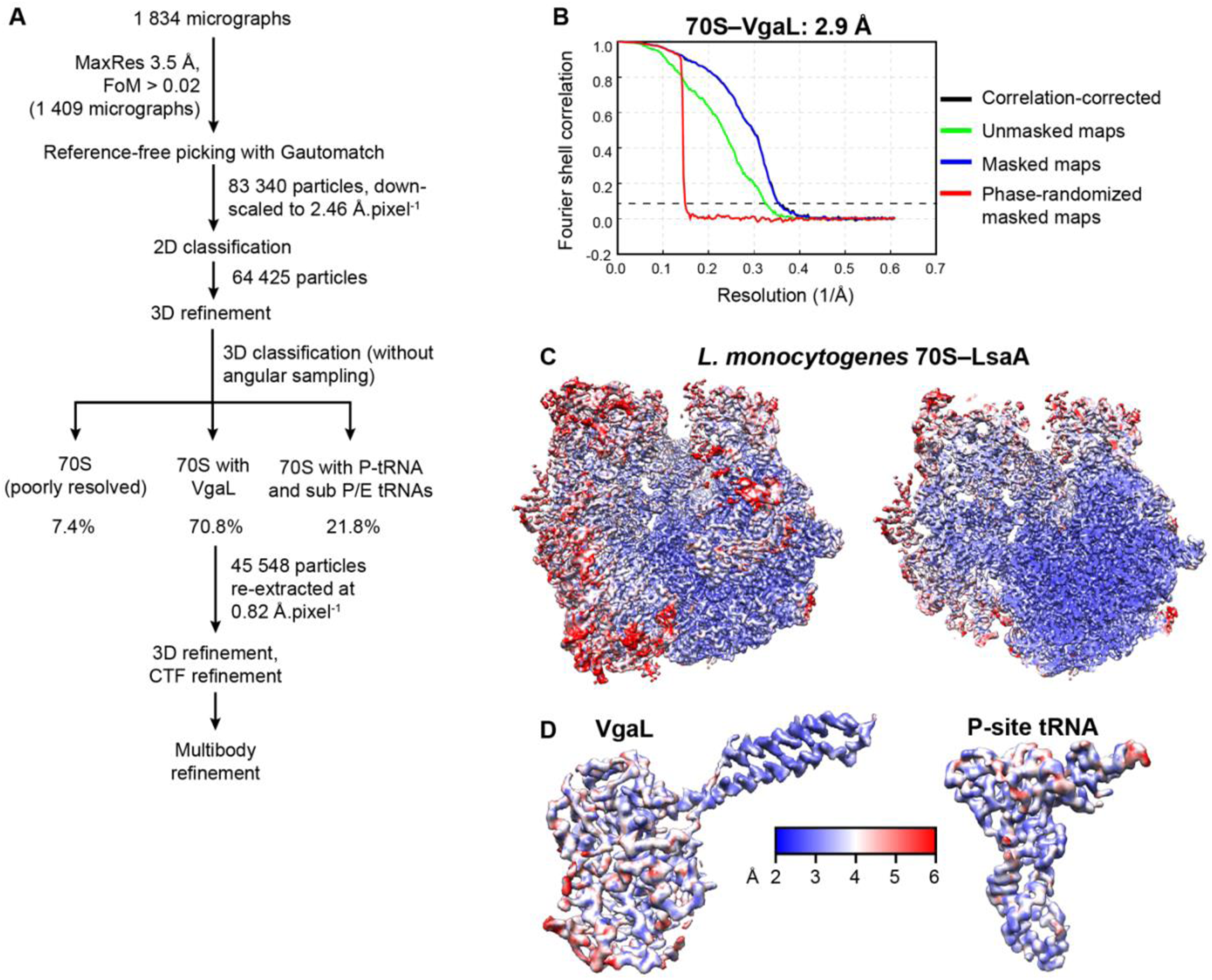
Processing of the cryo-EM data of VgaL–70S complex. **(A)** Processing scheme for the VgaL–70S complex. **(B)** Fourier Shell Correlation (FSC) curves of the VgaL-70S complex with a dashed line at 0.143 indicating average resolutions of 2.9 Å. **(C)** Overview (left) and transverse section (right) of the cryo-EM map of the VgaL–70S complex coloured according to local resolution. **(D)** Isolated density of VgaL (left) and P-site tRNA (right) from the VgaL-70S complex coloured according to local resolution.

**Figure S7.**
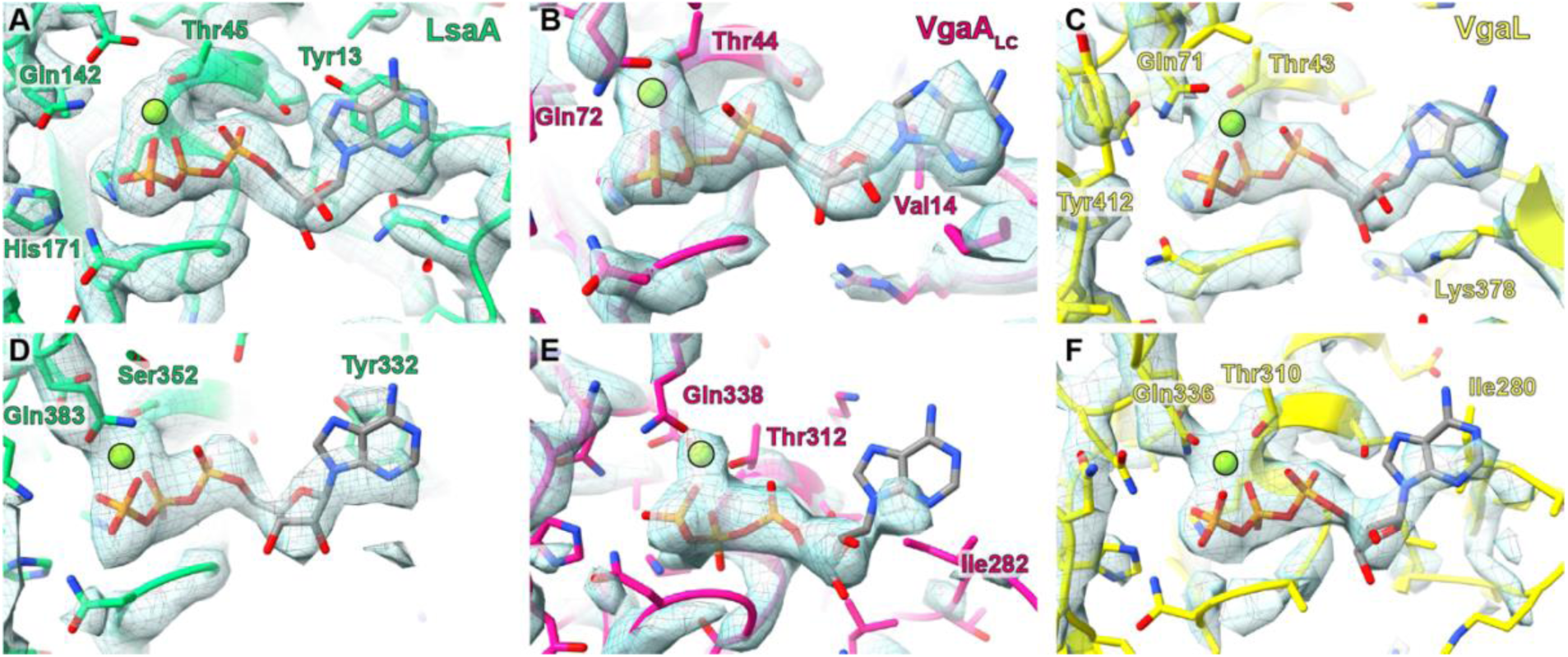
ATP in the ARE-bound 70S structures. Model and density surrounding the innermost ATP bound by LsaA **(A)**, VgaA_LC_ **(B)**, and VgaL **(C)** viewed from the direction of the signature sequence of NBD2 (model and density not shown). A black outline highlights a magnesium ion. **D–F**, as for **A–C** except for the peripheral nucleotide-binding site viewed from the direction of the signature sequence of NBD1 (model and density not shown). Density from post-processed maps is shown.

**Figure S8.**
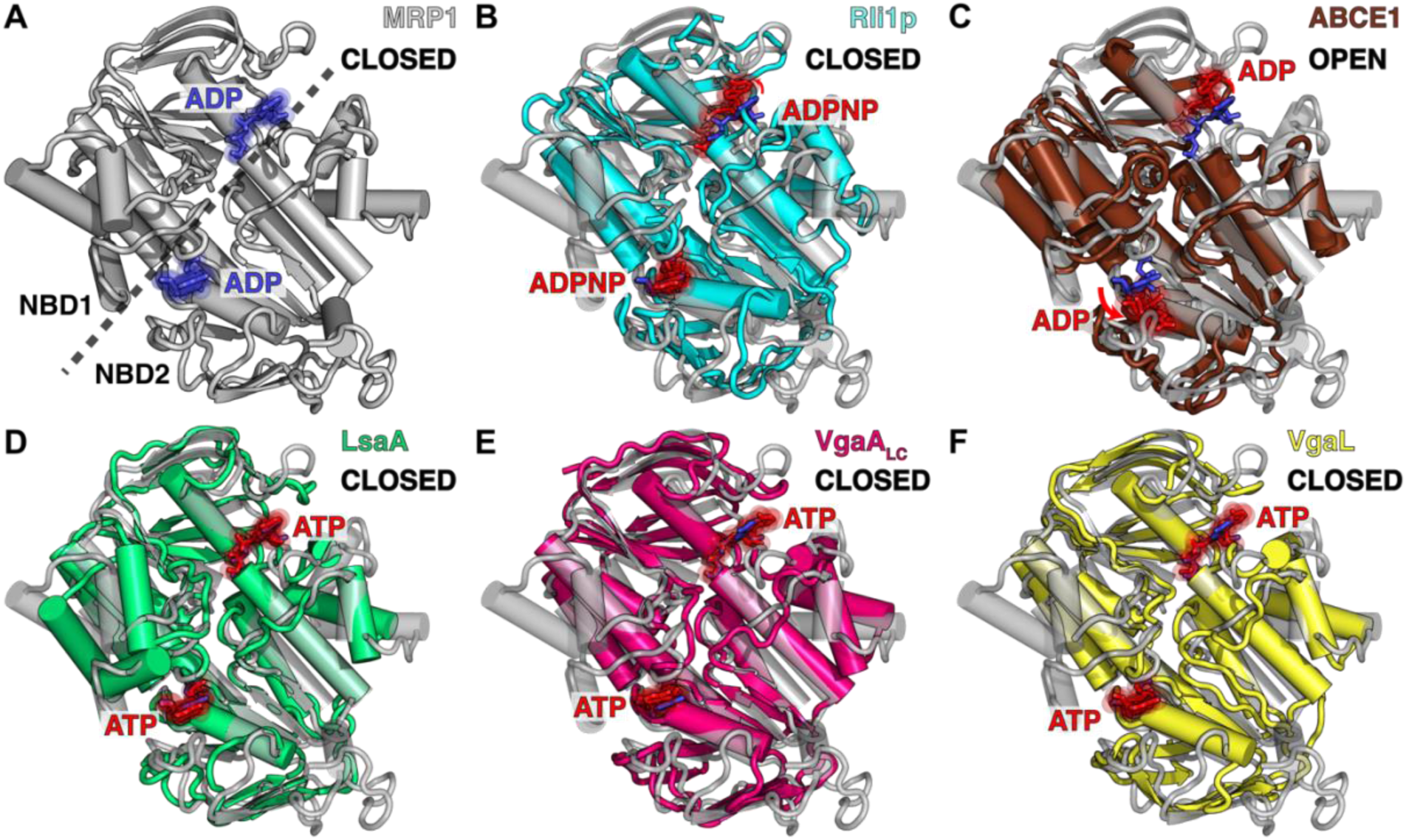
LsaA, VgaA_LC_ and VgaL NBDs exhibit a closed conformation. **(A)** The closed conformation of the multidrug transporter MRP1 (grey) with bound ADP molecules (blue, PDB 6BHU) (Johnson & Chen, 2018). **(B, C)** Alignment (based on NBD1) and superimposition of the closed conformation of MRP1 from **(A)** with the ABC domains of **(B)** Rli1p (cyan) in closed conformation with bound ADPNP (red, PDB 5LL6) (Heuer *et al*, 2017), **(C)** ABCE1 (brown) in open conformation with bound ADP (red, PDB 3J15) (Becker *et al*, 2012), and with (D–F) closed ARE ABCF NBD conformations with bound ATP (red) for **(D)** LsaA (green), **(E)** VgaA_LC_ (magenta) and **(F)** VgaL (yellow).

**Figure S9.**
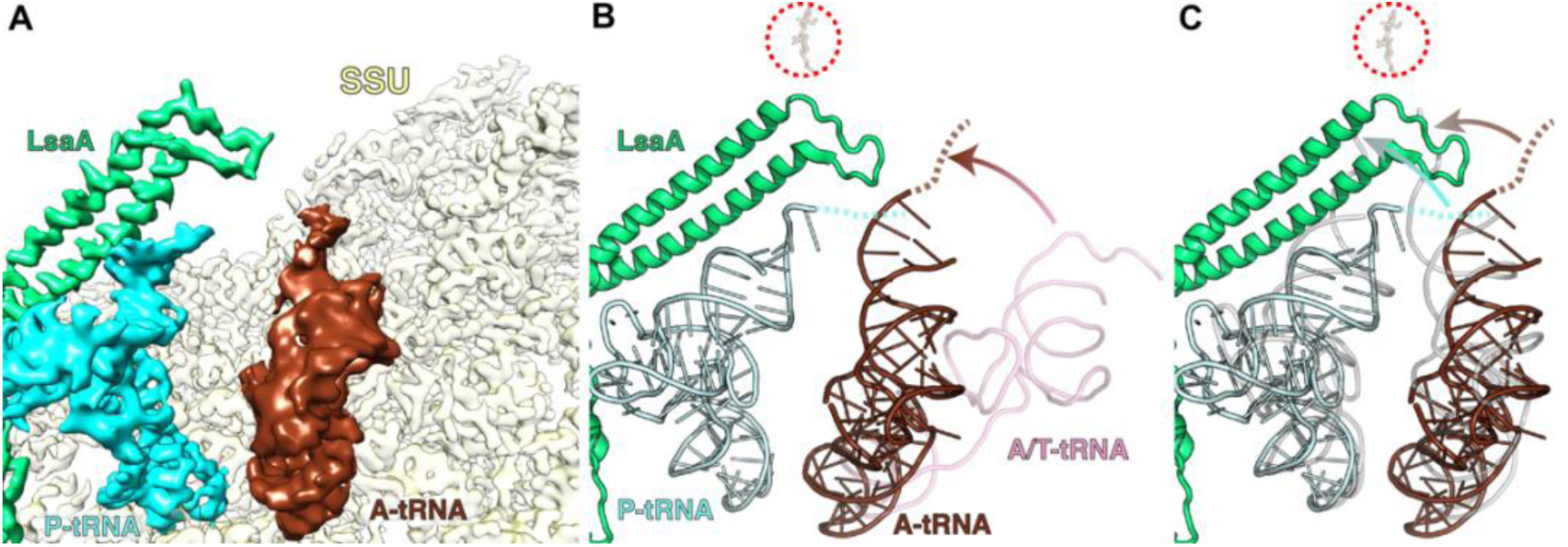
Presence of A-site tRNA in the LsaA-70S complex. **(A)** Cryo-EM map density for LsaA (green), P-site tRNA (cyan) and A-site tRNA (brown) in the LsaA–70S complex with A-site tRNA. Density for small subunit (yellow) is shown for reference. Density for the large subunit is not shown. **(B)** The same view as A, except with molecular models. The brown dashed line indicates a likely path for the 3′ CCA end of the distorted A-tRNA. A pre-accommodation A/T tRNA (pink, PDB 4V5L) (Voorhees *et al*, 2010) is superimposed. The position of the lincomycin binding site (red dotted circle) is shown for comparison (PDB 5HKV) (Matzov *et al*., 2017). **(C)** Similar to **(B)** except with classical accommodated A- and P-site tRNAs from pre-attack state superimposed (both grey, PDB 1VY4) (Polikanov *et al*., 2014).

**Figure S10.**
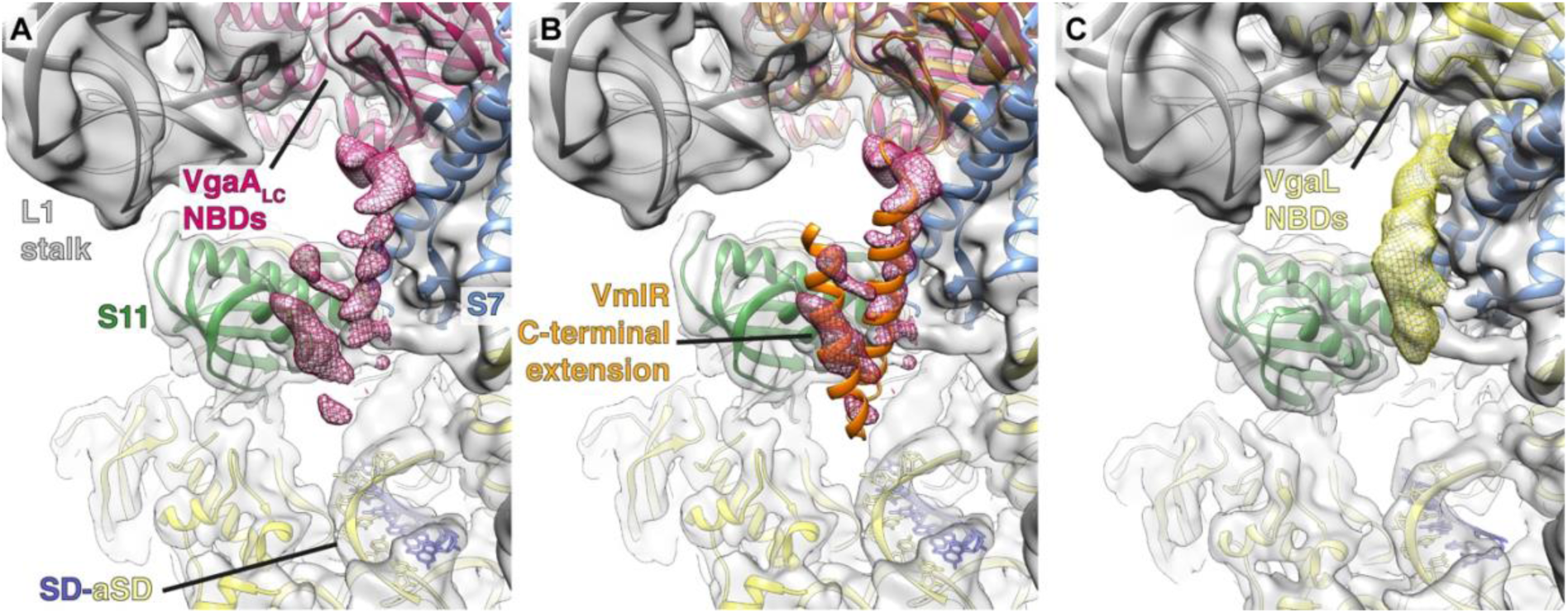
C-terminal extensions of VgaA_LC_ and VgaL on the small subunit. (**A-C**) Cryo-EM map (grey) with molecular model for (**B–C**) VgaA_LC_–70S complex, and (**C**) VgaL– 70S complex, showing density for L1 stalk (grey) on the large subunit, and ribosomal proteins S7 (blue), S11 (green) as well as the SD–anti-SD helix on the small subunit (yellow). In (**A**) and (**B**), density for the C-terminal extension (CTE) of VgaA_LC_ (magenta mesh) is fragmented, and in (**B**) fitted with the model of the CTE from VmlR (orange, PDB 6HA8) (Crowe-McAuliffe *et al*., 2018) based on alignment of the NBDs. In (**C**), density for the C-terminal extension (CTE) of VgaL (yellow mesh) also reaches between the S7–S11 cleft and is consistent with an α-helical conformation, but appears to be distinct from VmlR and VgaA_LC_ and could not be modelled at this resolution.

**Figure S11.**
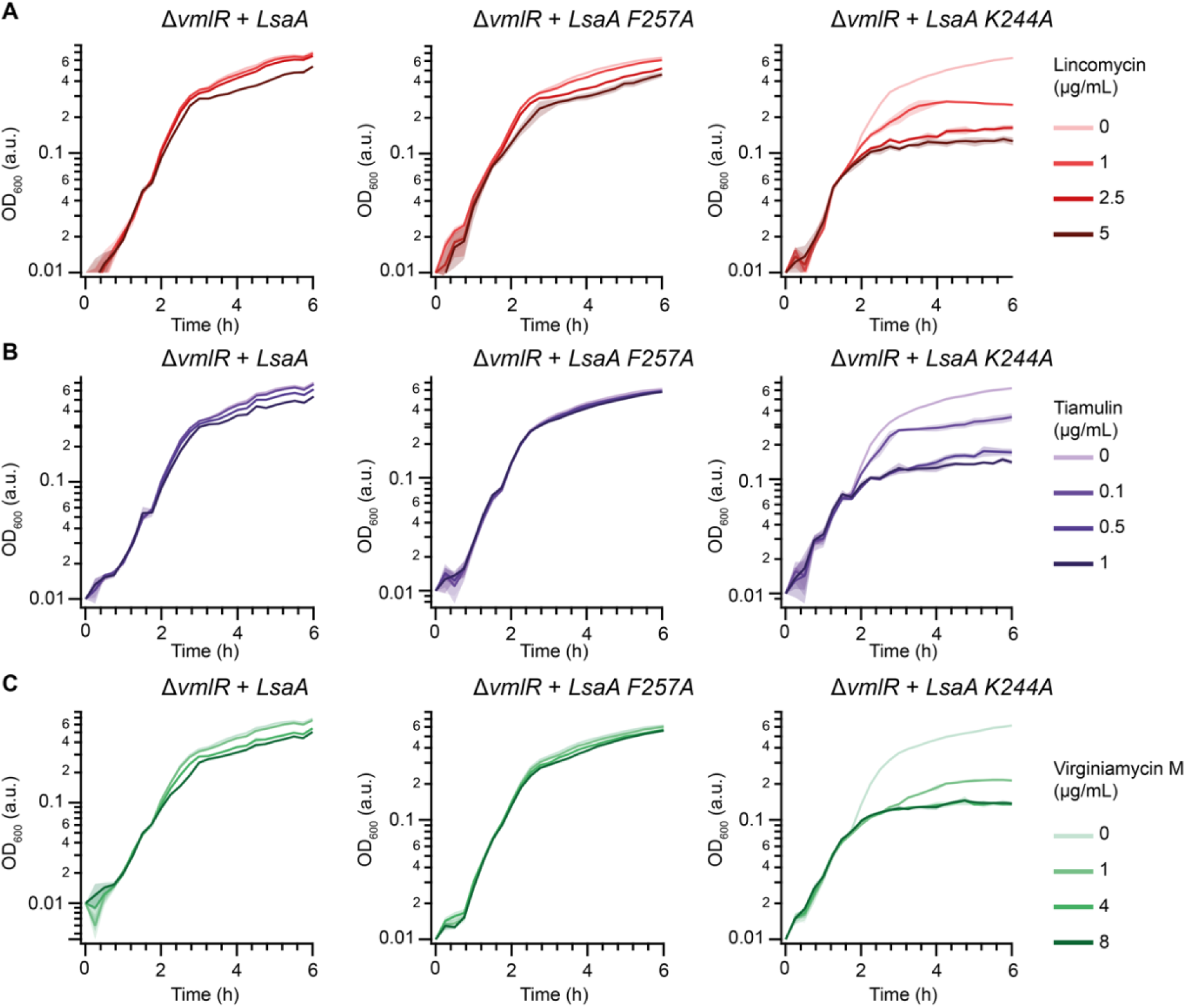
Effect of amino acid substitutions in ARD on antibiotic resistance in LsaA. Growth of *B. subtilis* Δ*vmlR* expressing the indicated LsaA variants over time in the presence of lincomycin (**A**), tiamulin (**B**), and virginiamycin M (**C**). *B. subtilis* strains (VHB109, 168 and 169) were grown in LB media with 1 mM IPTG at 37 °C with medium shaking. At the 90 minutes time point (OD_600_ ≍ 0.1) antibiotics were added to the final concentrations as indicated on the figure. The SD of three biological replicates is indicated with pale shading.

**Figure S12.**
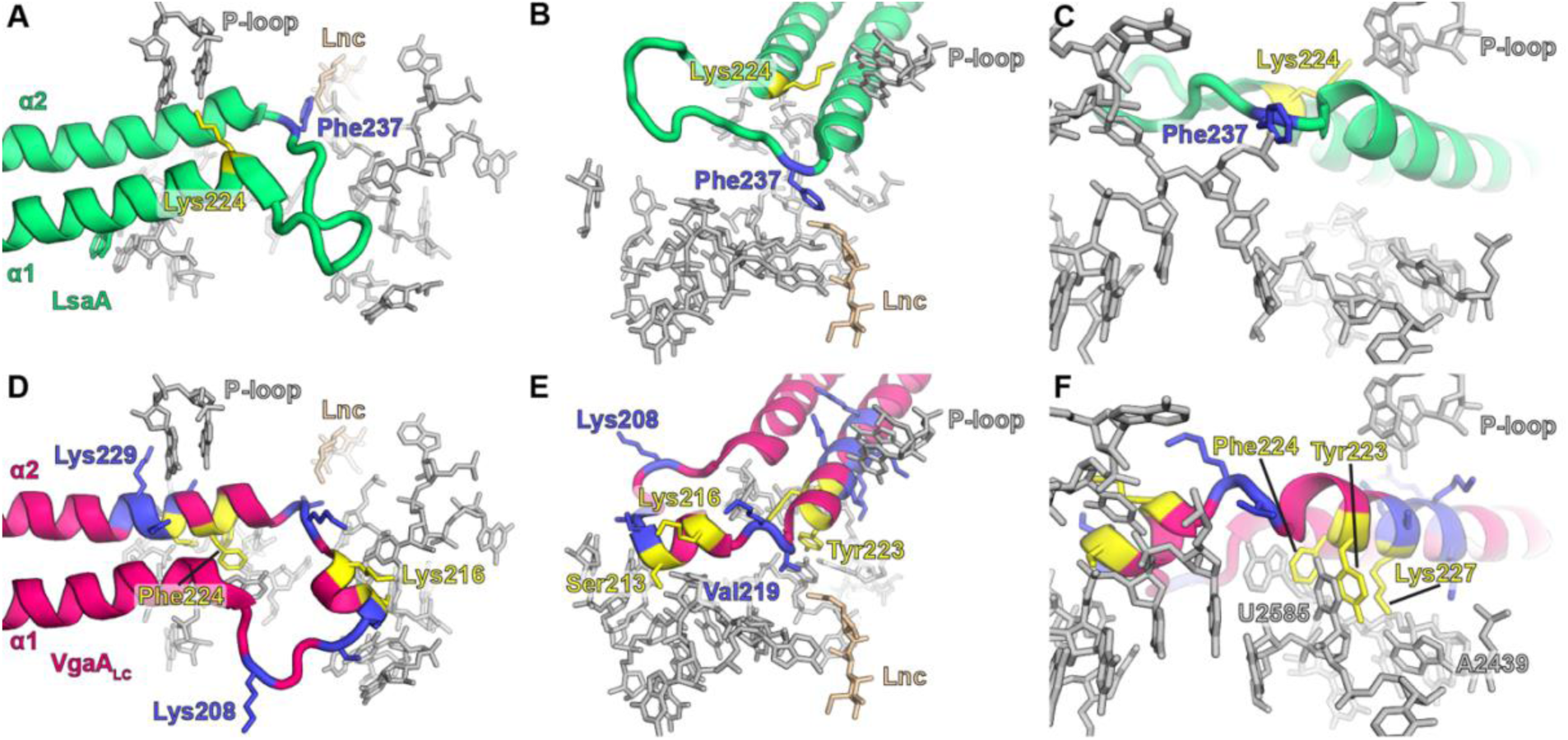
Visualisation of tested mutations in VgaA_LC_ and LsaA. Residues in blue did not affect antibiotic resistance when mutated to alanine, and residues in yellow reduced antibiotic resistance when mutated to alanine. **A–C**, three views of the LsaA ARD with selected *E. faecalis* 23S 23S rRNA nucleotides shown. **D–F**, three views of the VgaA_LC_ ARD with selected *S. aureus* 23S 23S rRNA nucleotides shown. See also Tables S1 and S2.

**Figure S13.**
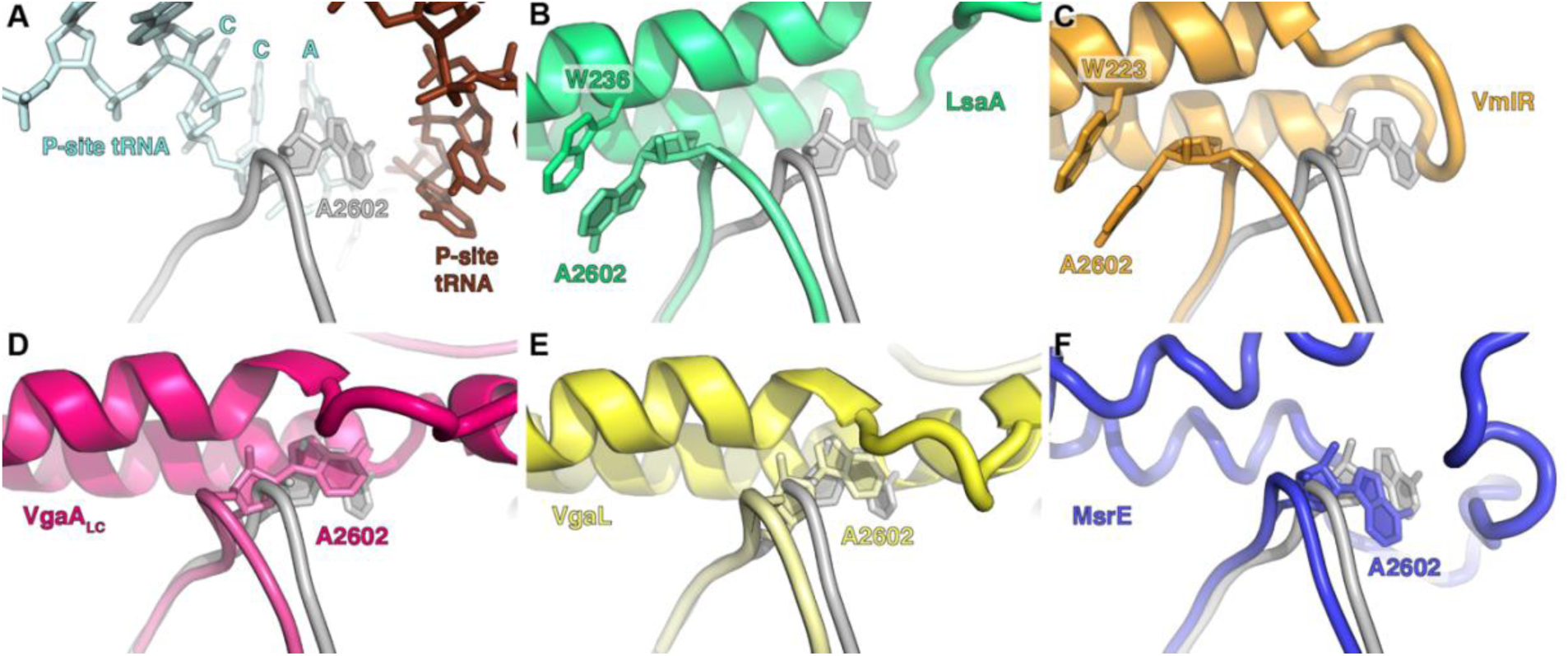
Comparison of A2602 position between ribosomes with and without bound AREs. **(A)** A2602 with accommodated A- and P-site tRNAs in the ‘pre-attack’ state (PDB 1VY4) (Polikanov *et al*., 2014) **(B)** Conformation of A2602 with bound LsaA with 23S rRNA from **(A)**. **(C–F)** Similar to **(B)**, except for VmlR, VgaA_LC_, VgaL, and MsrE (Crowe-McAuliffe *et al*., 2018; Su *et al*., 2018).

**Figure S14.**
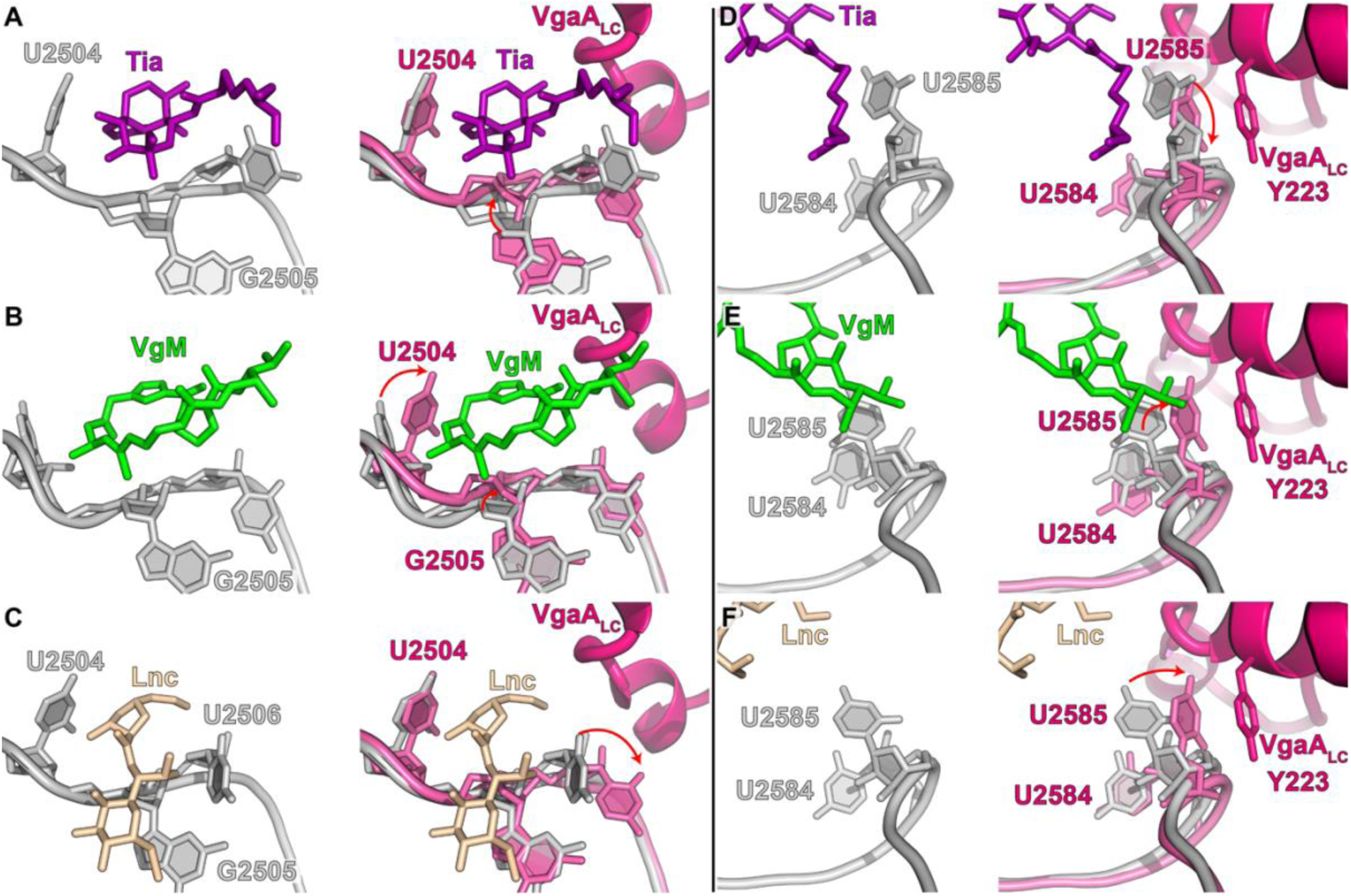
Conformation of the PTC in the presence of VgaA_LC_ and antibiotics. (**A–C**) The conformation of selected 23S rRNA nucleotides at the PTC in the presence of either (**A**) tiamulin (Tia, purple, PDB 1XBP) (Schlünzen *et al*., 2004), (**B**) virginiamycin M (VgM, green, PDB 1YIT) (Tu *et al*., 2005), or (**C**) lincomycin (Lnc, tan, PDB 5HKV) (Matzov *et al*., 2017). Left panels show the antibiotic-bound structures only, right panels have superimposed nucleotides and protein from the VgaA_LC_-bound ribosome (pink). (**D–F**) As for A–C, except with focus on U2585. Red arrows indicate significant shifts in nucleotide positions from antibiotic-bound to VgaA_LC_-bound ribosomes.

**Figure S15.**
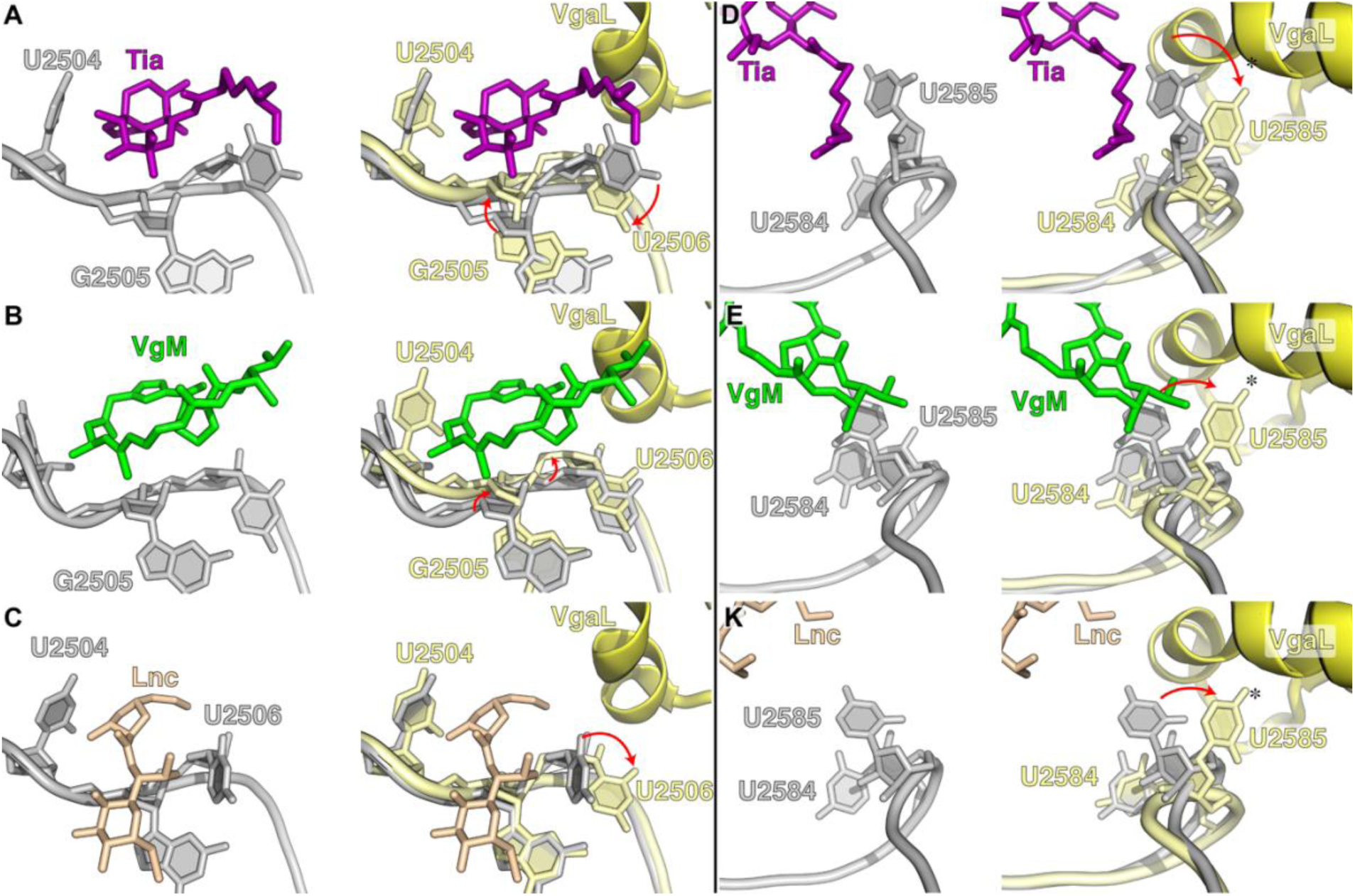
Conformation of the PTC in the presence of VgaL and antibiotics. The conformation of selected 23S rRNA nucleotides at the PTC in the presence of either (**A**) tiamulin (Tia, purple, PDB 1XBP) (Schlünzen *et al*., 2004), (**B**) virginiamycin M (VgM, green, PDB 1YIT) (Tu *et al*., 2005), or (**C**) lincomycin (Lnc, tan, PDB 5HKV) (Matzov *et al*., 2017). Left panels show the antibiotic-bound structures only, right panels have superimposed nucleotides and protein from the VgaL-bound ribosome (yellow). (**D–F**) As for **A**–**C**, except with focus on U2585. Red arrows indicate significant shifts in nucleotide positions from the antibiotic-bound to VgaL-bound ribosome. An asterisk indicates low confidence in the position of U2585 due to weak density.

**Figure S16.**
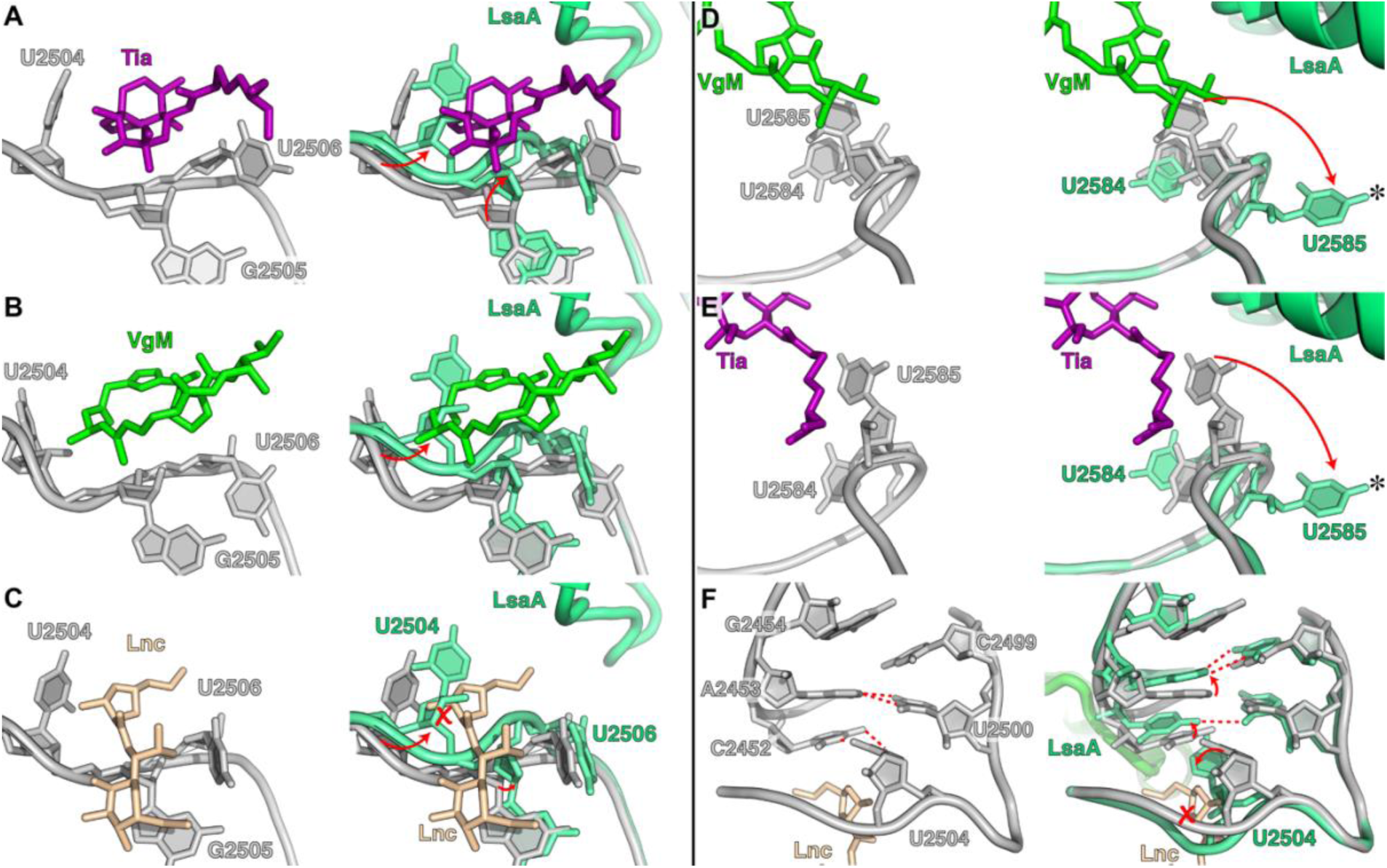
Conformation of the PTC in the presence of LsaA and antibiotics. (**A–C**) The conformation of selected 23S rRNA nucleotides at the PTC in the presence of either (**A**) tiamulin (Tia, purple, PDB 1XBP) (Schlünzen *et al*., 2004), (**B**) virginiamycin M (VgM, green, PDB 1YIT) (Tu *et al*., 2005), or (**C**) lincomycin (Lnc, tan, PDB 5HKV) (Matzov *et al*., 2017). Left panels show the antibiotic-bound structures only, right panels have superimposed nucleotides and protein from the LsaA-bound ribosome (green). (**D–F**) As for A–C, except with focus on U2585 (**D**, **E**) or U2504 (**F**). Red arrows indicate significant shifts in nucleotide positions from antibiotic-bound to LsaA-bound ribosomes, and red crosses indicate significant overlap between the lincomycin-binding site and U2504 in the LsaA-bound ribosome. An asterisk indicates low confidence in the position of U2585 due to weak density.

**Table S1.** Minimum inhibitory concentrations (MICs) of ribosome-targeting antibiotics against E. faecalis expressing LsaA. 5 x 10^5^ CFU/mL (OD_600_ approximately 0.0005) of either *E. faecalis* OG1RF, Δ*lsaA* (*lsa::Kan*) strain TX5332 transformed with empty pCIE_spec_ plasmid, or with pCIE_spec_ derivative for expression of LsaA was used to inoculate BHI media supplemented with 2 mg/mL kanamycin to prevent *lsa* revertants, 0.1 mg/mL spectinomycin to maintain the pCIE_spec_ plasmid, 100 ng/mL of cCF10 peptide to induce expression of LsaA as well as increasing concentrations of antibiotics. After 16-20 hours at 37 °C without shaking, the presence or absence of bacterial growth was scored by eye. The MIC values that exceed the empty vector control are shown in bold.

| antibiotic class | antibiotic | MIC, µg/mL |  |  |  |
| --- | --- | --- | --- | --- | --- |
|  |  | *E. faecalis*<br>OG1RF | *E. faecalis*<br>TX5332<br>pCIE<sub>spec</sub><br>(VHp426) | *E. faecalis*<br>TX5332<br>pCIE<sub>spec</sub> *lsaA*<br>(VHp431) | *E. faecalis*<br>TX5332<br>pCIE *lsaA*-<br>*HTF* |
| phenicols | chloramphenicol | 2-4 | 2-4 | 2-4 |  |
|  | thiamphenicol | 4 | 4 | 4 |  |
|  | florfenicol | 1 | 1-2 | 1-2 |  |
| oxazolidinones | linezolid | 1 | 1 | 1 | 1 |
| macrolides | erythromycin | 1 | 0.5-1 | 0.5 | 0.5 |
|  | azithromycin | 1-2 | 0.5-1 | 0.5-1 |  |
|  | leucomycin | 0.5-1 | 0.5 | 0.5-1 |  |
| lincosamides | lincomycin | **32** | 0.125 | **16-32** | **8-16** |
|  | clindamycin | **16-32** | 0.0156 | **16** | **4-8** |
| pleuromutilins | tiamulin | **128** | 0.0625 | **128** | **32-64** |
|  | retapamulin | **>64** | 0.0156 | **>64** |  |
| streptogramins | virginiamycin M1 | **>64** | 4 | **>128** |  |
|  | virginiamycin S1 | 8 | 8 | 8 |  |
| tetracyclines | tetracycline | 0.5 | 0.25 | 0.25 |  |

**Table S2.** Minimum inhibitory concentrations (MICs) of ribosome-targeting antibiotics against S. aureus expressing VgaA_LC_. S. aureus strain SH1000, harbouring empty vector pRMC2 or pRMC2 expressing wild-type *vgaA_LC_* or its mutants.

| Construct<br>(mutation) | MIC, µg/mL |  |  |  |  |
| --- | --- | --- | --- | --- | --- |
|  | lincomycin | clindamycin | tiamulin | retapamulin | virginiamycin M1 |
| pRMC2 | 0.5 | 0.06 | 0.5 | 0.06 | 2 |
| pRMC2: *vgaA<sub>LC</sub>* | **16** | **2** | **8** | **4** | 4 |
| pRMC2: *vgaA<sub>LC</sub>*<br>(K<sub>208A</sub>) | **16** | **2** | **16** | **8** | 4 |
| pRMC2: *vgaA<sub>LC</sub>*<br>(S<sub>211A</sub>) | **16** | **2** | **16** | **4** | 4 |
| pRMC2: *vgaA<sub>LC</sub>*<br>(S<sub>212A</sub>) | **8** | **2** | **16** | **8** | 4 |
| pRMC2: *vgaA<sub>LC</sub>*<br>(S<sub>213A</sub>) | **2** | 0.125 | 1 | 0.125 | 1 |
| pRMC2: *vgaA<sub>LC</sub>*<br>(K<sub>216A</sub>) | **8** | **0.5** | **4** | **1** | 4 |
| pRMC: *vgaA<sub>LC</sub>*<br>(K<sub>218A</sub>) | **16** | **1** | **16** | **4** | 4 |
| pRMC2: *vgaA<sub>LC</sub>*<br>(V<sub>219A</sub>) | **16** | **1** | **16** | **8** | 2 |
| pRMC2: *vgaA<sub>LC</sub>*<br>(W<sub>223A</sub>) | **2** | 0.125 | 1 | 0.125 | 1 |
| pRMC2: *vgaA<sub>LC</sub>*<br>(F<sub>224A</sub>) | 0.5 | 0.06 | 0.25 | 0.06 | 1 |
| pRMC2: *vgaA<sub>LC</sub>*<br>(S<sub>226A</sub>) | **16** | **2** | **16** | **8** | 4 |
| pRMC2: *vgaA<sub>LC</sub>*<br>(K<sub>227A</sub>) | **4** | **0.25** | 1 | 0.125 | 2 |
| pRMC2: *vgaA<sub>LC</sub>*<br>(G<sub>228A</sub>) | **8** | **2** | **8** | **2** | 2 |
| pRMC2: *vgaA<sub>LC</sub>*<br>(K<sub>229A</sub>) | **16** | **2** | **8** | **4** | 4 |
| pRMC2: *vgaA<sub>LC</sub>*<br>(K<sub>230A</sub>) | **16** | **2** | **8** | **4** | 4 |
| pRMC2: *vgaA<sub>LC</sub>*<br>(R<sub>232A</sub>) | **16** | **2** | **8** | **2** | 4 |

**Table S3.**
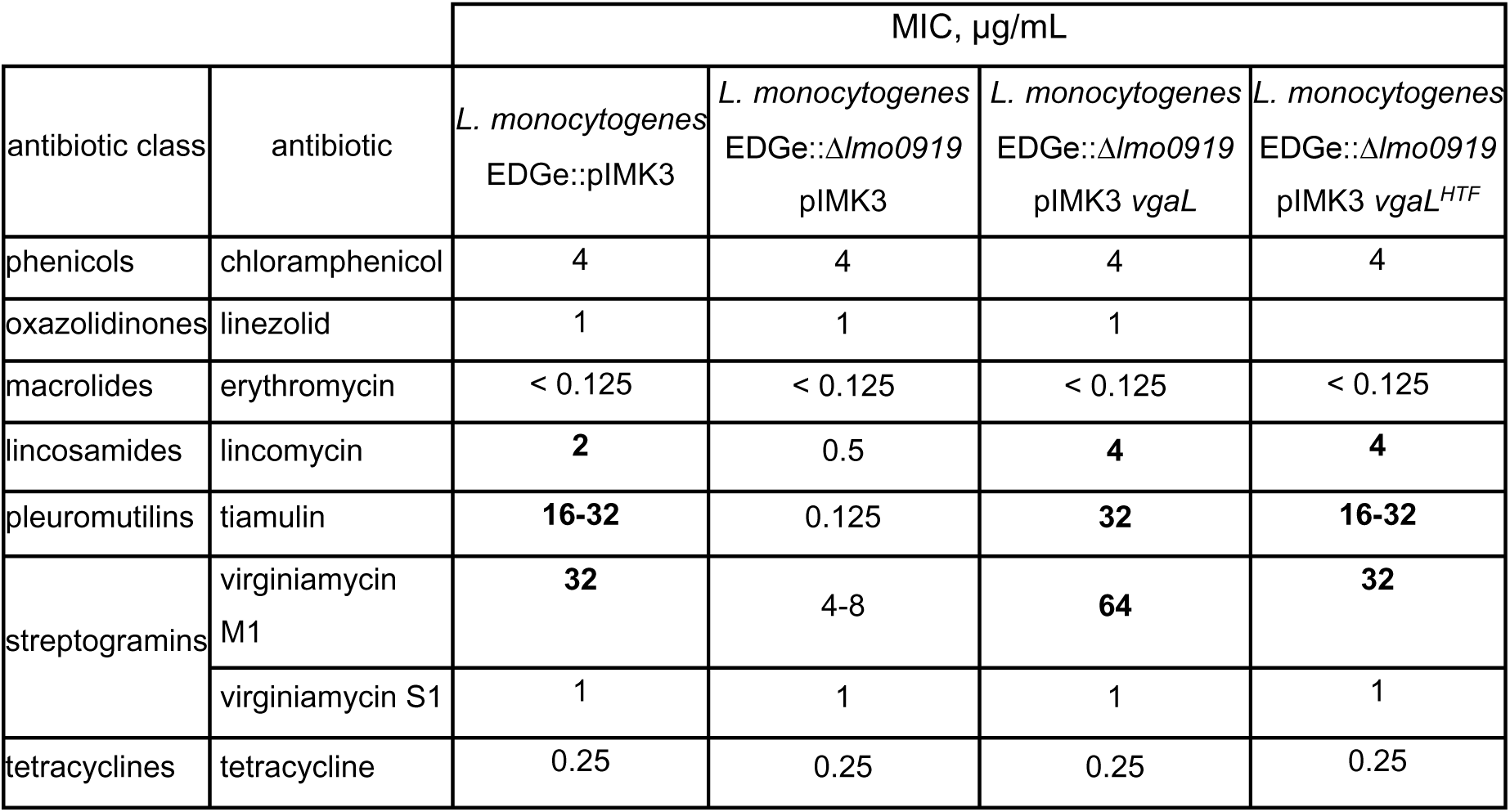
Minimum inhibitory concentrations (MICs) of ribosome-targeting antibiotics against *L. monocytogenes* EDG-*e* expressing VgaL (Lmo0919). 5 x 10^5^ CFU/mL (approximately OD_600_ 0.0003) of *L. monocytogenes EDGe*, Δ*lmo0919* (markerless) strain with integrated empty pIMK3 plasmid, or with pIMK3 encoding VgaL or VgaL-HTF was used to inoculate BHI media supplemented with 50 µg/mL kanamycin to maintain the integrative pIMK3 plasmid, 1 mM IPTG to induce expression of VgaL as well as increasing concentrations of antibiotics. After 16-20 hours at 37 °C without shaking, the presence or absence of bacterial growth was scored by eye. The MIC values that exceed the empty vector control lacking chromosomal *lmo0919* are shown in bold.

| antibiotic class | antibiotic | MIC, µg/mL |  |  |  |
| --- | --- | --- | --- | --- | --- |
| | | *L. monocytogenes*<br>EDGe::pIMK3 | *L. monocytogenes*<br>EDGe::Δ*lmo0919*<br>pIMK3 | *L. monocytogenes*<br>EDGe::Δ*lmo0919*<br>pIMK3 *vgaL* | *L. monocytogenes*<br>EDGe::Δ*lmo0919*<br>pIMK3 *vgaL*<sup>HTF</sup> |
| phenicols | chloramphenicol | 4 | 4 | 4 | 4 |
| oxazolidinones | linezolid | 1 | 1 | 1 |  |
| macrolides | erythromycin | < 0.125 | < 0.125 | < 0.125 | < 0.125 |
| lincosamides | lincomycin | **2** | 0.5 | **4** | **4** |
| pleuromutilins | tiamulin | **16-32** | 0.125 | **32** | **16-32** |
| streptogramins | virginiamycin M1 | **32** | 4-8 | **64** | **32** |
|  | virginiamycin S1 | 1 | 1 | 1 | 1 |
| tetracyclines | tetracycline | 0.25 | 0.25 | 0.25 | 0.25 |

**Table S4.** Cryo-EM data collection, modelling and refinement statistics.

|  | *E. faecalis*<br>70S–LsaA | *S. aureus*<br>70S–P-tRNA | *S. aureus*<br>70S–VgaA<sub>Lc</sub> | *L. monocytogenes*<br>70S–VgaL |
| --- | --- | --- | --- | --- |
| **Data collection** |  |  |  |  |
| Magnification (×) | 130 000 | 165 000 | 165 000 | 165 000 |
| Electron dose (e-/Å<sup>2</sup>) | 38.0 | 26.3 | 26.3 | 28.28 |
| Defocus range (μm) | −0.7–2.2 | −0.7–1.9 | −0.7–1.9 | −0.8–2.0 |
| Pixel size (Å) | 1.041 | 0.82 | 0.82 | 0.82 |
| Initial particles | 61 009 | 165 827 | 165 827 | 83 340 |
| Final particles | 59 262 | 61 910 | 35 129 | 45 548 |
| Average resolution (Å) | 2.9 | 3.1 | 3.1 | 2.9 |
| **Model composition** |  |  |  |  |
| Atoms | 144 982 | 139 909 | 145 651 | 144 492 |
| Protein residues | 5 753 | 5 330 | 5 783 | 5 715 |
| RNA bases | 4 627 | 4554* | 4 647 | 4617 |
| **Refinement** |  |  |  |  |
| Map CC around atoms | 0.85 | 0.89 | 0.88 | 0.86 |
| Map CC whole unit cell | 0.85 | 0.88 | 0.87 | 0.85 |
| Map sharpening B | −35.42 | −56.43 | −62.31 | −68.16 |
| R.M.S. deviations |  |  |  |  |
| Bond lengths (Å) | 0.008 | 0.010 | 0.009 | 0.014 |
| Bond angles (°) | 0.906 | 0.947 | 0.933 | 1.082 |
| **Validation** |  |  |  |  |
| MolProbity score | 1.74 | 1.73 | 1.76 | 1.75 |
| Clash score | 4.19 | 3.87 | 4.24 | 4.41 |
| Poor rotamers (%) | 0.02 | 0.07 | 0.06 | 0.04 |
| Ramachandran |  |  |  |  |
| Favoured (%) | 90.48 | 89.88 | 89.64 | 90.76 |
| Outlier (%) | 0.16 | 0.02 | 0.05 | 0.02 |

\*23S rRNA helices H76–H78 of the L1 stalk were flexible and not modelled.

**Table S5.** Strains and Plasmids used in this study. Plasmid and strain construction is described in detail in supplemental text. *Denotes a plasmid constructed by the PEP facility at Umeå University.

| Strain | Description | Source |
| --- | --- | --- |
| *L. monocytogenes* EDGe | Wild-type serotype 1/2a strain | (Glaser *et al.*, 2001) |
| *L. monocytogenes* EDGe::pIMK3 | EGDe with empty pIMK3 plasmid containing P<sub>help</sub> promoter integrated at tRNA<sup>Arg</sup> locus | This work |
| *L. monocytogenes* EDGe::pIMK3/*lmo0919*<sup>HTF</sup> | EGDe with VgaL-HTF overexpressed from the P<sub>help</sub> promoter integrated at tRNA<sup>Arg</sup> locus | This work |
| *L. monocytogenes* EDGe::pIMK3/*lmo0919*<sup>EQ2-HTF</sup> | EGDe with VgaL EQ2-HTF overexpressed from the P<sub>help</sub> promoter integrated at tRNA<sup>Arg</sup> locus | This work |
| *L. monocytogenes* EDGe::Δ*lmo0919* | EGDe harboring a *lmo0919* marker less deletion lacking VgaL | This work |
| *L. monocytogenes* EDGe::Δ*lmo0919*::pIMK3 | EGDe::Δ*lmo0919* with empty pIMK3 plasmid containing P<sub>help</sub> promoter integrated at tRNA<sup>Arg</sup> locus | This work |
| *L. monocytogenes* EDGe::Δ*lmo0919*::pIMK3/*lmo0919* | EGDe::Δ*lmo0919* with VgaL overexpressed from the P<sub>help</sub> promoter integrated at tRNA<sup>Arg</sup> locus | This work |
| *L. monocytogenes* EDGe::Δ*lmo0919*::pIMK3/*lmo0919*<sup>HTF</sup> | EGDe::Δ*lmo0919* with VgaL-HTF overexpressed from the P<sub>help</sub> promoter integrated at tRNA<sup>Arg</sup> locus | This work |
| *L. monocytogenes* EDGe::Δ*lmo0919*::pIMK3/*lmo0919*<sup>HTF-EQ2</sup> | EGDe::Δ*lmo0919* with VgaL-EQ2-HTF overexpressed from the P<sub>help</sub> promoter integrated at tRNA<sup>Arg</sup> locus | This work |
| *E. faecalis* OG1RF | Rif<sup>r</sup> Fus<sup>r</sup>; WT *E. faecalis* | (Singh *et al.*, 2002) |
| *E. faecalis* TX5332 | Rif<sup>r</sup> Fus<sup>r</sup> Kan<sup>r</sup>; *lsa* gene disruption mutant (OG1RF *lsa*::pTEX4577) | (Davis *et al.*, 2001) |
| *S. aureus* SH1000 | Functional *rsbU*<sup>+</sup> derivative of *S. aureus* 8325-4 | (Horsburgh *et al.*, 2002; O'Neill, 2010) |
| *E. coli* S17.1 | *E. coli* strain used for conjugative plasmid transfer to *L. monocytogenes* | (Simon *et al.*, 1983) |

| Plasmid | Description | Reference |
| --- | --- | --- |
| pIMK3 | Kan<sup>r</sup>; Listerial tRNA<sup>Arg</sup> locus specific integrative vector for high-level IPTG-induced protein expression from the P<sub>help</sub> promoter | (Monk *et al.*, 2008) |
| pMAD | Amp<sup>r</sup>, Ery<sup>r</sup>; *lacZ*; thermosensitive shuttle vector used for allelic exchange in *L. monocytogenes* | (Arnaud *et al.*, 2004) |
| pHT009 | Amp<sup>r</sup>, Km<sup>r</sup>; thrC locus specific integrative vector for high-level IPTG-induced protein expression from the P<sub>hy-spnak</sub> promoter | (Crowe-McAuliffe *et al.*, 2018) |
| VHp689 | pMAD Δ*lmo0919* | This work |
| VHp690 | pIMK3:*lmo0919* | This work |
| VHp692 | pIMK3:*lmo0919-HTF* | This work |
| VHp693 | pIMK3:*lmo0919-EQ2-HTF* | This work |
| pTX5333 | Cm<sup>r</sup>; *E. faecalis*-*E. coli* shuttle plasmid expressing LsaA from native promoter | (Singh *et al.*, 2002) |
| pCIE | Cm<sup>r</sup>; *E. faecalis*-*E. coli* shuttle plasmid for cCF10 induced expression of proteins | (Weaver *et al.*, 2017) |
| VHp100 | pCIE:*lsaA-HTF* | This work* |
| VHp149 | pCIE:*lsaA-EQ2-HTF* | This work* |
| VHp369 | pHT009-*lsaA* | This work |
| VHp426 | pCIE, Sc<sup>r</sup>; Cm<sup>r</sup> gene swapped to spectinomycin resistance (Sc<sup>r</sup>) gene | This work* |
| VHp431 | VHp426:*lsa* | This work* |
| VHp526 | pHT009-*lsaAK244A* | This work |
| VHp526 | pHT009-*lsaAK244A* | This work |
| pRMC2 | Amp<sup>r</sup>, Cm<sup>r</sup>; *E. coli* - *S. aureus* shuttle plasmid for tetracycline-regulable expression of proteins in the latter host. | (Corrigan & Foster, 2009) |
| pRMC2:*vgaA-FLAG<sub>3</sub>* | pRMC2 expressing C-terminally FLAG<sub>3</sub> tagged VgaA<sub>LC</sub> | This work |
| pRMC2:*vgaA-EQ2-FLAG<sub>3</sub>* | pRMC2 expressing C-terminally FLAG<sub>3</sub> tagged VgaA<sub>LC</sub>-E<sub>105Q</sub>, E<sub>410Q</sub> | This work |
| pRMC2:*vgaA<sub>LC</sub>* | pRMC2 expressing wild-type VgaA<sub>LC</sub> | This work |
| pRMC2:*vgaA<sub>LC</sub>* (K<sub>208A</sub>) | pRMC2 expressing VgaA<sub>LC</sub><sup>K208A</sup> | This work |
| pRMC2:*vgaA<sub>LC</sub>* (S<sub>211A</sub>) | pRMC2 expressing VgaA<sub>LC</sub><sup>S211A</sup> | This work |
| pRMC2:*vgaA<sub>LC</sub>* (S<sub>212A</sub>) | pRMC2 expressing VgaA<sub>LC</sub><sup>S212A</sup> | This work |
| pRMC2:*vgaA<sub>LC</sub>* (S<sub>213A</sub>) | pRMC2 expressing VgaA<sub>LC</sub><sup>S213A</sup> | This work |
| pRMC2:*vgaA<sub>LC</sub>* (K<sub>216A</sub>) | pRMC2 expressing VgaA<sub>LC</sub><sup>K216A</sup> | This work |
| pRMC2:*vgaA<sub>LC</sub>* (K<sub>218A</sub>) | pRMC2 expressing VgaA<sub>LC</sub><sup>K218A</sup> | This work |
| pRMC2:*vgaA<sub>LC</sub>* (V<sub>219A</sub>) | pRMC2 expressing VgaA<sub>LC</sub><sup>V219A</sup> | This work |
| pRMC2:*vgaA<sub>LC</sub>* (W<sub>223A</sub>) | pRMC2 expressing VgaA<sub>LC</sub><sup>W223A</sup> | This work |
| pRMC2:*vgaA<sub>LC</sub>* (F<sub>224A</sub>) | pRMC2 expressing VgaA<sub>LC</sub><sup>F224A</sup> | This work |
| pRMC2:*vgaA<sub>LC</sub>* (S<sub>226A</sub>) | pRMC2 expressing VgaA<sub>LC</sub><sup>S226A</sup> | This work |
| pRMC2:*vgaA<sub>LC</sub>* (K<sub>227A</sub>) | pRMC2 expressing VgaA<sub>LC</sub><sup>K227A</sup> | This work |
| pRMC2:*vgaA<sub>LC</sub>* (G<sub>228A</sub>) | pRMC2 expressing VgaA<sub>LC</sub><sup>G228A</sup> | This work |
| **pRMC2: *vgaA<sub>LC</sub>* (K<sub>229</sub>A)** | pRMC2 expressing VgaA<sub>LC</sub><sup>K229A</sup> | This work |
| **pRMC2: *vgaA<sub>LC</sub>* (K<sub>230</sub>A)** | pRMC2 expressing VgaA<sub>LC</sub><sup>K230A</sup> | This work |
| **pRMC2: *vgaA<sub>LC</sub>* (A<sub>232</sub>A)** | pRMC2 expressing VgaA<sub>LC</sub><sup>A232A</sup> | This work |

## Notes

### Competing Interest Statement

The authors have declared no competing interest.

